# Bidirectional modulation of human emotional conflict resolution using intracranial stimulation

**DOI:** 10.1101/825893

**Authors:** Angelique C. Paulk, Ali Yousefi, Kristen K. Ellard, Kara Farnes, Noam Peled, Britni Crocker, Rina Zelmann, Deborah Vallejo-Lopez, Gavin Belok, Sam Zorowitz, Ishita Basu, Afsana Afzal, Anna Gilmour, Daniel S. Weisholtz, G. Reese Cosgrove, Bernard S. Chang, Jeffrey E. Arle, Ziv M. Williams, Uri T. Eden, Thilo Deckersbach, Darin D. Dougherty, Emad N. Eskandar, Alik S. Widge, Sydney S. Cash

## Abstract

The ability to regulate emotions in the service of meeting ongoing goals and task demands is a key aspect of adaptive human behavior in our volatile social world. Consequently, difficulties in processing and responding to emotional stimuli underlie many psychiatric diseases ranging from depression to anxiety, the common thread being effects on behavior. Behavior, which is made up of shifting, difficult to measure hidden states such as attention and emotion reactivity, is a product of integrating external input and latent mental processes. Directly measuring, and differentiating, separable hidden cognitive, emotional, and attentional states contributing to emotion conflict resolution, however, is challenging, particularly when only using task-relevant behavioral measures such as reaction time. State-space representations are a powerful method for investigating hidden states underlying complex systems. Using state-space modeling of behavior, we identified relevant hidden cognitive states and predicted behavior in a standardized emotion regulation task. After identifying and validating models which best fit the behavior and narrowing our focus to one model, we used targeted intracranial stimulation of the emotion regulation-relevant neurocircuitry, including prefrontal structures and the amygdala, to causally modulate separable states. Finally, we focused on this one validated state-space model to perform real-time, bidirectional closed-loop adaptive stimulation in a subset of participants. These approaches enable an improved understanding of how to sample and understand emotional processing in a way which could be leveraged in neuromodulatory therapy for disorders of emotional regulation.

## Introduction

To function in a complex world, humans must continuously regulate responses to conflicting emotional cues in the service of meeting ongoing, goal-directed demands. Defined as emotion regulation, humans have the ability to adaptively influence which emotions are experienced, when they are experienced, and how they are expressed given a current context (1–4). The process of emotion regulation has been well studied over the past 20 years (1, 3, 5–9) and it is clear that adaptive emotion regulation requires an ability to detect the salience of an internally or externally generated emotional cue, and to subsequently attend towards or away from this cue and adjust behavior accordingly dependent upon the current situational demands. Deficits in emotion regulation have been shown to play a role and contribute to symptom severity across anxiety, mood, depression, PTSD, and related disorders (1–4, 9–18). Thus, emotion dysregulation represents a key transdiagnostic dimension of psychiatric disease and a target for intervention. At the level of neurocircuitry, emotion regulation recruits a network of cortical regions implicated in executive control functions (e.g. attention orienting, working memory) including dorsal and ventrolateral prefrontal cortex (dlPFC, vlPFC), dorsomedial PFC (dmPFC) and rostral and dorsal anterior cingulate cortex (rACC, dACC; (18–28)). Increased activation in these regions is associated with decreased activation in subcortical limbic regions including amygdala (29–31). Thus, successful emotion regulation increases executive control in the service of goal directed activities while subsequently decreasing salience processing (26, 28–31).

The development of novel neurotherapeutics to target emotion dysregulation requires a precise understanding of the relationship between activation along emotion regulation-related neurocircuitry and behavior (32–35). Successful emotion regulation likely involves a large number of hidden cognitive dynamics that ultimately contribute to behavioral responses, including shifts in attention, cognitive flexibility, emotion reactivity, and behavioral adaptability (1, 5). Therefore, understanding how regions within the identified neural circuitry of emotion regulation enhance or inhibit these hidden cognitive states is crucial to the identification of viable and directed targets for intervention using neurotherapeutics, including invasive and non-invasive neuromodulation (34, 36–38). Recent promising advancements in identifying states of mood relative to neural activity have led to the use of closed loop tools to stimulate based on these neural states, with stimulation in the orbitofrontal cortex resulting in the improvement of mood (35). However, mood is the product of a numerous underlying hidden states and processes including emotion regulation (14, 39). To develop the therapy further for focused relief shaped by underlying mechanisms, we propose identifying the underlying dynamics to plan for more targeted, and tailored, approaches supporting the overall modulation of emotion regulation.

State-space modeling is a powerful computational tool that can allow for the identification and examination of hidden features underlying behavior relevant to emotion regulation, including attention, cognitive flexibility, emotion reactivity, and adaptability. State-space modelling involves arriving at an estimated probabilistic dependence between the latent state variable, such as hidden features, and an observed measurement, such as reaction time or accuracy. This comes with the main assumption that behavior such as reaction time or accuracy is driven by a combination of hidden features which can be modelled as multivariate, latent cognitive states. In state space modelling, these latent states can vary over time, exhibit inherent dynamics and vary with external inputs, such as visual stimuli or trial type. An advantage of state-space modeling is that each feature can be modeled as a hidden cognitive state and estimated on a per trial basis using an expectation maximization approach derived from the behavior (40–42). Furthermore, since the state space modeling approach can be used to factorize the underlying cognitive components and estimate their dynamics over the course of experiment one can test a series of state-space models with different features and validate which best corresponds to behavior. This approach represents a significant improvement over multi-trial block-averaged approaches, wherein behavioral responses are examined cross-sectionally over entire task duration (e.g. >50-100 trials), thus losing potentially important information about the specific, nuanced states underlying behavior. Further, state-space models may allow for more precise mapping of behavior to neural activity, which enables more precise identification of targets for neurostimulation. The power of this state-space modeling approach has been shown in both learning and tests of cognitive flexibility (40, 43–47).

In the current study, we used a state space approach to capture specific hidden features of emotion regulation relevant to behavior during performance on the Emotion Conflict Resolution task (ECR; (6)), a well validated behavioral probe of emotion regulation that has been shown to induce activation of emotion regulation neurocircuitry (6, 22, 23). Indeed, ECR performance and brain activation during the task has been shown to predict drug treatment responsiveness in depression (48). An important point is that emotion is a key part of the ECR task since it requires identifying the emotion on a presented face while ignoring the overlaid word, requiring emotion perception and the activation of emotion circuitry (6). Yet, since emotion regulation, at its core, not only involves emotion perception but resolving emotion conflict to regulate emotion responses (5, 10), we chose the the ECR task since it addresses both emotion perception and resolving emotion conflict. By focusing on ECR, we are addressing one aspect of implicit emotion regulation, namely the ability to maintain goal-directed behaviors in the presence of competing/conflicting affective information. With the notion that there are several factors, including latent cognitive states, affecting behavioral responses to the ECR task at any point in time, thereby introducing unexplained ‘noise’ to the behavioral measure, we hypothesized that a state-space modeling approach would provide a more tractable measure that would allow access to the underlying, or hidden, cognitive states driving reaction time, such as attention or conflict resolution, on a per-trial basis. As such, performance on the ECR, as it is a modified Stroop task (49), requires a number of processes relevant to emotion regulation including salience processing (emotion reactivity), goal-directed processing (attentional control, response inhibition), recognition of a conflict between salience processing and goal-directed processing, resolution of this conflict through behavioral adaptation (cognitive flexibility and adaptability), and working memory (6, 19–23). As the task includes both difficult and easy trials, and the complexity of the responses, which include speed (reaction time) and accuracy, there could be numerous hidden cognitive dynamics contributing to behavioral responses. To validate whether these hidden cognitive dynamics could be identified and then separately altered through neuromodulation, we performed a sequence of steps. First, we applied a state-space approach purely to behavior (reaction time and accuracy) during the ECR task to model hidden states and to use these models to predict reaction time from three separate cohorts (healthy volunteers, psychiatric patients, and patients with intractable epilepsy). Second, after establishing a subset of behavioral models which were high-performing, in separate task sessions, we tested whether identified hidden states within these behavioral state space models can be driven by stimulation in specific brain networks. We hypothesized that direct electrical intracranial stimulation in different brain regions would have differential, and causal, effects on these behavioral features as has been hinted at with stimulation in other brain regions in learning and memory (50, 51), mood (35, 52), and OCD (36, 53, 54). As a final test, we used the state-space model in closed-loop adaptive stimulation to modulate behavior predictably.

## RESULTS

### Behavior in the Emotion Conflict Resolution task

To understand the hidden cognitive state underlying emotion conflict resolution and emotion reactivity, we examined the behavioral dynamics of three different groups of individuals while they performed the Emotion Conflict Resolution (ECR) task (6): 1) Healthy control volunteers (N=42); 2) individuals with psychiatric diagnoses (N=16) and 3) participants with intractable epilepsy undergoing intracranial recordings (N=41; **Supplemental Table 1**). During the task, participants identified the emotion of a face while ignoring an overlaid word that was either congruent (C) or incongruent (I) with the face’s emotion ((6); Fig. 1A, Supplemental Figure 1). Consistent with prior literature and across the three groups, incongruent trials induced longer reaction times (RTs) compared to congruent trials, and overall accuracy was above 88% ((6, 48); z-scored relative to all trials per participant, p<0.0001; Kruskal-Wallis test; Fig. 1B, **Supplemental Figure 1A**). Valence (Happy vs fear) did not significantly affect overall behavior (**Supplemental Fig. 2**).

**Fig. 1.**
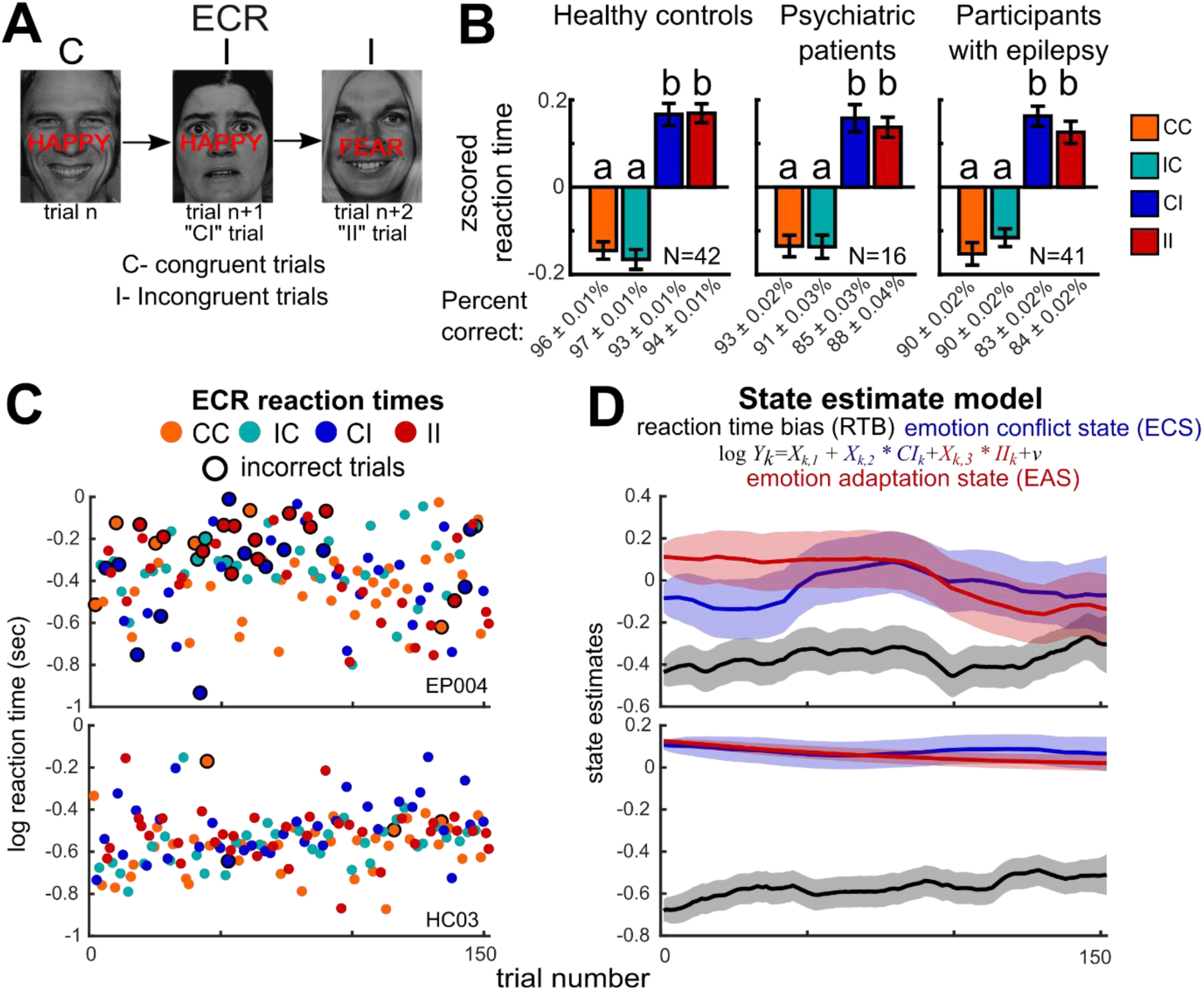
Emotion Conflict Resolution (ECR) task behavior as a model of human emotion regulation. **A. Task. B.** Participants respond to the emotion of the face while ignoring the word, which can match (congruent trials, C) or conflict (incongruent trials, I). Trials preceded by congruent or incongruent trials (e.g. CI, IC, II, and CC) induced different reaction times across the patient groups which included healthy control volunteers, participants diagnosed with psychiatric disorders, and participants with epilepsy. All reaction times were z-scored relative to the all trials per task run (RTs). The data set (N=99) included 42 healthy controls, 16 participants diagnosed with a psychiatric disorder, and 41 participants with epilepsy. Z-scored RTs are significantly different between trial types per group (p<0.00001, Kruskal-Wallis test). **C.** The reaction times of two participants (EP004 and HC003) during the task demonstrating reaction time variability across trials**. D**. The ECR state estimate the Conflict-Adaptation Model and its three main components: the overall reaction time state (Bias), the emotion conflict state (CI), and the adaptation state (II). State estimates during the task (top, participant EP004, bottom, HC03), with the confidence bounds highlighted as shaded regions.

### Self-reported emotion reactivity and emotion regulation

Self-report psychometric questionnaires were used as an independent measure of emotion reactivity, anxiety and emotion regulation to the central ECR task. Significant differences in responses to self-report psychometric scales between the three participant groups were found across three measures of emotion reactivity and regulation ability (Emotion Reactivity Scale, ERS; Difficulties in Emotion Regulation Scale, DERS; Anxiety Sensitivity Index, ASI; See **Methods**; see **Supplemental Table 2** for details of average raw scale scores by participant group; **Supplemental Fig. 2;** (8, 55–57)). Consistent with the existing literature, relative to healthy controls, individuals diagnosed with psychiatric disorders endorsed significantly greater emotion reactivity (*p*=0.00001; Wilcoxon rank sum; ERS), and difficulties with emotion regulation (*p*=0.0003; Wilcoxon rank sum; DERS), but not anxiety sensitivity after correcting for multiple comparisons (*p*=0.032; Wilcoxon rank sum; ASI; **Supplemental Fig. 2**). Patients with epilepsy had responses which were widely distributed, overlapping between the other two groups (**Supplemental Fig. 2**) with scores not significantly different to either the healthy control group or the patients diagnosed with psychiatric disorders following corrections for multiple comparisons (p>0.0245; Wilcoxon rank sum), consistent with the known co-morbidity of epilepsy and psychiatric symptoms (58). No significant correlation was found between z-scored scale scores and block-averaged reaction time performance on the ECR task (average zscored RT difference between congruent to incongruent trials minus incongruent to incongruent trials (CI-II), or accuracy).

### State-space modelling of conflict and adaptation behavior in the ECR task

We developed and validated behavioral state estimate model(s) which best describe latent variables underlying reaction times and accuracy using only non-stimulated behavioral task sessions (see **Methods; Supplemental Fig. 3-4**). The approach involved modelling hidden (latent) cognitive states from the trial by trial behavior (e.g. RT) with the assumption that these states are driven both by their underlying dynamics and exogenous input such as trial type. To do this, we assumed the hidden states are influenced by, and therefore have a linear relationship with, indicator terms such as trial type or trial history (40, 41). Estimating those hidden states required an expectation maximization approach with multiple iterations per task session as well as estimations of noise inherent to the system (see **Methods**). In addressing the hidden states using state-space approaches, we could regress out unrelated components or RT or state-related changes in RT using state space approaches, thereby identifying, and measuring, features of cognitive flexibility, attention, and conflict adaptation or conflict resolution. The methodology involved identifying state space models which addressed known task-related cognitive processes such as cognitive flexibility or reaction time bias, testing how well these models fit the non-stimulated behavioral data, how well each model predicted reaction times, and their correspondence to psychometric measures of emotion reactivity, as compared to a healthy control group (8, 55–57) (**Supplemental Fig. 3-4**). The choice of the models tested, including what trial types (e.g. whether a trial was congruent or incongruent, trial history, etc.) and whether we used reaction time, accuracy, or both, to model, and predict, reaction time was informed by the large body of literature regarding the underlying hidden cognitive states driving behavior during ECR and, more generally, Stroop tasks ((6, 19–21, 23); see **Methods**). Each model was used to address an idea from the literature regarding Stroop tasks, such as the number of trials or trial difficulty, and whether the hidden cognitive states indeed varied over time (see **Methods**; **Supplemental Fig. 3-4**). Interestingly, when we included trial accuracy either alone or modeled along with reaction time in a mixed effects model, the models either did not converge or had very large noise terms, indicating models including accuracy did not fit, or predict, the reaction time or accuracy as well as a reaction time-only modelling approach (see **Methods**). Ultimately, eight of thirty models survived the criteria testing (**Supplemental Fig. 3-4**). All eight models had three features in common distinguishing them from the other criteria: 1) All viable models had a reaction time bias term which addresses the overall drift in reaction time over the task and 2) 5 of the 8 models had a transition term relating to whether the task trial type switched from congruent to incongruent (CI) or vice versa (incongruent to congruent, IC). We decided to focus on two of these eight viable models since these two models (Model 1:the Conflict-Adaptation model and Model 15: the Conflict-Adaptation equipoise model) included terms which relate to resolving surprising emotion conflict (as when transitioning between an easy to difficult trial, e.g. congruent to incongruent trial, CI) and emotion adaptation (such as when adapting to increased difficulty following a previous difficult trial, e.g. incongruent trial to incongruent trial, II; (6, 22, 48); Fig. 1). The emotion conflict (CI) and adaptation (II) terms have been proposed to relate to emotion regulation (6, 23). Thus, the Conflict-Adaptation model (specifically Conflict minus Adaptation; Fig. 1E; 2A, **see Methods**) reflects the idea that adaptation to a pattern of two difficult trials in a row (Incongruent trial to Incongruent trial, II) can be measured as an emotion adaptation state (EAS), whereas the switch burden of Congruent trial to Incongruent trial (CI) is measured as the emotion conflict state (ECS; (6, 23, 48)). One problem with Model 1 is that we had to estimate ECS and EAS separately and subtract the mean ECS-EAS terms (e.g. Conflict-Adaptation) to derive a trial by trial term describing the balance of surprise (Conflict) and adapting to difficulty (Adaptation). For this reason, we developed Model 15, the Conflict-Adaptation equipoise model, which is a reformulation that allowed us to measure the Conflict-Adaptation balance, or equipoise, as a single hidden state variable with confidence bounds and associated noise and confidence metrics (see **Methods**).

**Figure 2.**
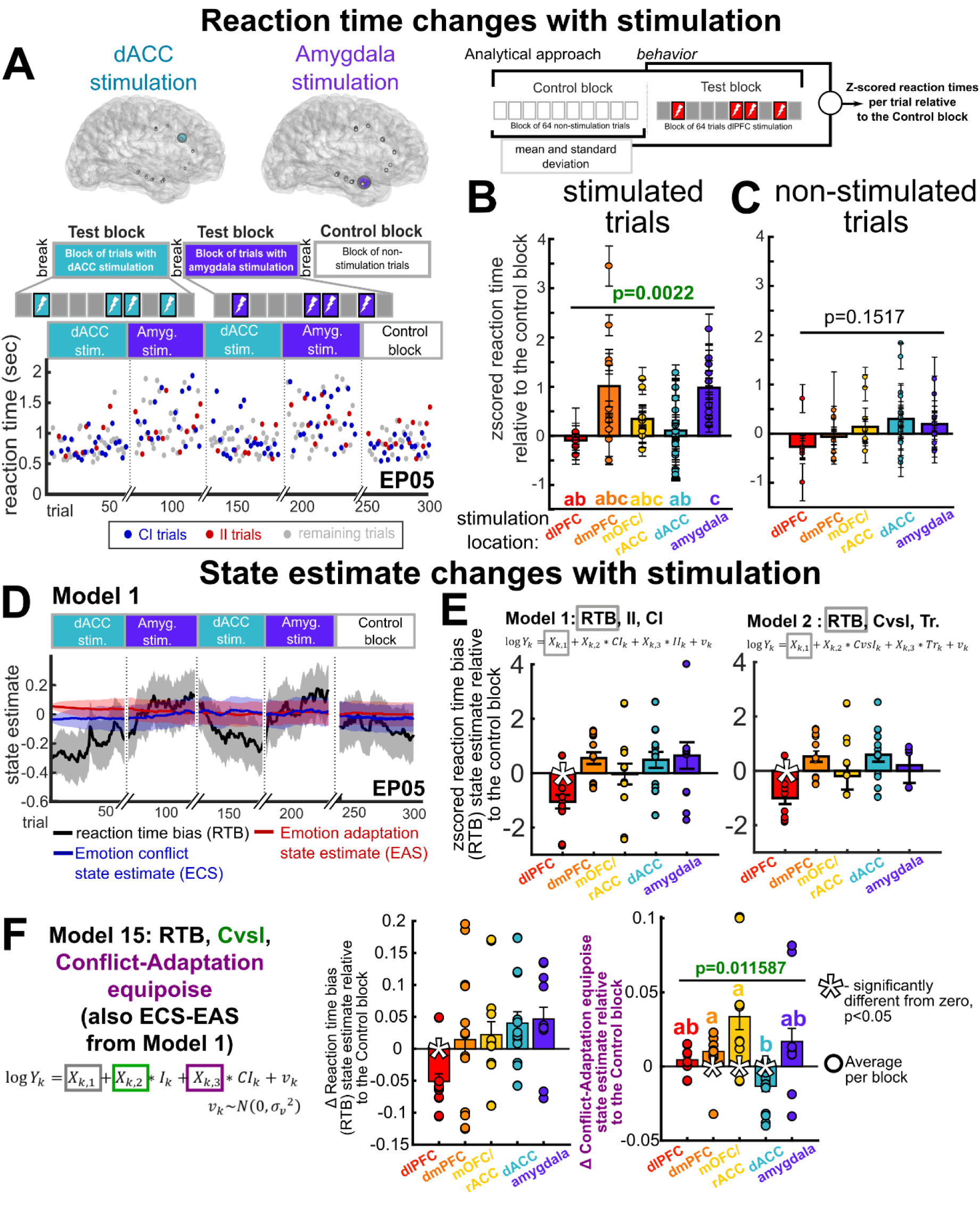
Neural stimulation induces behavioral changes in reaction time and conflict-adaptation state. **A**. RTs during Test and Control blocks with dACC and amygdala stimulation (participant EP05). X-axis breaks: ~8-minute pauses. **B**. Z-scored RTs were significantly different between stimulation sites during stimulated trials (p=0.0022; Kruskal-Wallis test; Chi-Sq.=31). **C**. The non-stimulated trials during Test blocks had no significant changes in zscored RTs (p=0.1517; Kruskal-Wallis test) **D**. ECR state estimate Conflict-Adaptation (Model 1) with three main components: the overall reaction time bias (Bias, RTB), the emotion conflict state (ECS, CI), and the adaptation state (EAS, II). RTB shifts with amygdala and dACC stimulation (95% confidence bounds indicated in shaded areas). **E**. RTB state estimate changes for both Models 1 and 2, particularly during dlPFC stimulation. **F-G.** Both the Conflict-Adaptation state (**G**) and EAS (**F**) varied significantly across stimulation sites (N=13; p>0.05; Kruskal-Wallis test). (the letters a-b along the x axis indicate statistically separable groups, *post hoc* Tukey-Kramer testing; *-p<0.0033, significantly different from zero, Wilcoxon sign rank test). In **B** and **F**, error bars indicate standard error across trials. Abbreviations: Dorsolateral (dl) and dorsomedial (dm) prefrontal cortex-dlPFC, dmPFC; medial (m) orbitofrontal cortex – mOFC; dorsal (d) and rostral (r) anterior cingulate-dACC, rACC.

#### Modeling the emotion conflict (ECS) and adaptation (EAS) states

For the Conflict-Adaptation model (Model 1), maximum likelihood (ML) estimates over 1000 model iterations showed stable model convergence (**Supplemental Figure 4)**, low average model noise (*V*=0.0317±0.0196; **Supplemental Figure 4**) and low average state variable noise (*W*<0.00601), suggesting good model fit to the reaction time behavior. Further, the model demonstrated good predictive ability with the squared difference between predicted and actual log reaction times low (squared difference=0.05±0.031). Furthermore, the reaction time bias term in the Conflict-Adaptation model was significantly positively correlated with scores on psychometric measures of anxiety (ERS: *r = 0.2900, p= 0.1271;* DERS: *r = 0.1343*, *p* = 0.4957; **ASI: *r = 0.4452*, *p = 0.0155***). In addition, the emotion conflict state (ECS; CI) was negatively correlated with scores on psychometric measures, significantly only for scores of emotion regulation (ERS: *r = −0.2576, p= 0.1773;* DERS: ***r = −0.4205, p = 0.0259***; ASI: *r = −0.1881*, *p = 0.3286*).

#### Modeling Conflict-Adaptation equipoise

For the Conflict-Adaptation equipoise model (Model 15), maximum likelihood (ML) estimates over 1000 model iterations again showed stable model convergence (**Supplemental Figure 4)**, low average model noise (*V=*0.0318±0.0196; Fig. 2A, E, F) and low average state variable noise (*W*<0.00604), suggesting good model fit to the reaction time behavior. Further, the model demonstrated good predictive ability (squared difference=0.05±0.031). The reaction time bias term in the conflict-avoidance model was significantly associated with scores on psychometric measures of anxiety (ERS: *r = 0.2987, p= 0.1155;* DERS: *r = 0.1386*, *p* = 0.4819; **ASI: *r = 0.4471*, *p = 0.0150***).

In sum, we were able to identify a subset of models which both fit the ECR task reaction time behavior for a large data set of individuals which were interpretable, did not over-fit the data, which mapped somewhat to psychometric questionnaires, and which allowed us to presumptively understand the hidden states underlying emotion conflict resolution. Narrowing our focus primarily to two models, namely the Conflict-Adaptation and the Conflict-Adaptation equipoise models, we then examined whether features of the models could be modulated with direct electrical stimulation during a separate set of task sessions performed by the participants with intractable epilepsy with implanted electrodes.

### Direct electrical stimulation can bidirectionally alter reaction time

To examine whether stimulation in brain regions known to be involved in emotion regulation and engaged during the ECR task (22, 48) results in reaction time and accuracy changes, we performed targeted stimulation in dACC, rACC, amygdala, dmPFC, and dlPFC, during performance in the ECR task (stimulation locations in **Supplemental Figure 6**). During Test blocks, non-stimulation trials were interspersed with stimulated trials (Fig. 2A). Stimulation in dmPFC, rACC and amygdala significantly altered RTs (dmPFC: *p=0.0391*; rACC: *p=0.0098*; amygdala: *p=0.002*; Wilcoxon signed rank test; Fig. 2B) but did not change accuracy (*all p-values >0.31*; Wilcoxon rank-sum). Stimulation of dmPFC, rACC, and amygdala increased RTs across all trial types (Fig. 2B), while stimulation in the dlPFC decreased RT (though not significant) and dACC stimulation produced split effects (Fig. 2B). Across regions, the biggest difference in z-scored RT was between the amygdala and the dlPFC stimulation, confirming existing literature on the dissociable roles these two brain regions may have during emotion regulation, processing, and reappraisal (26–29, 31). These effects were focal in time: z-scored reaction times during non-stimulated trials of stimulation blocks were not significantly different trials regardless of stimulation location during stimulated trials (*p=0.1517; Chi-Sq.= 6.7149*; Kruskal-Wallis test; Fig. 2C).

### Neural stimulation causally alters hidden cognitive states

These results point to a brain region specific focal effect of stimulation on ECR task behavior. However, this analysis reported average changes in reaction time regardless of trial type or trial number and did not address separate hidden cognitive states. As we had identified hidden cognitive states in a subset of viable state space models, we hypothesized we could differentially modulate these states using neural stimulation. If these state space models capture a hidden cognitive state, then it stood to reason that stimulation in certain brain regions could causally induce network changes resulting in behavioral changes corresponding with these altered hidden cognitive states. We, therefore, applied the state-space models to data obtained during stimulation experiments and measured the changes in the main components each model. Overall, across models, stimulation of the dlPFC resulted in significantly decreased reaction time bias relative to non-stimulation blocks in the same task session (p<.05, Wilcoxon signed rank test). This corresponded to the general decrease in reaction time (Fig. 2), a measure of attentional or effortful drift. The dlPFC stimulation effect on reaction time bias (RTB) was evident even in multiple viable behavioral state space models, including Model 2 which only included RTB, the current trial congruence (CvsI) or whether the trial was different to the previous trial (Tr., see **Methods**; Fig. 2E). In contrast, dACC stimulation resulted in significantly increased reaction time bias across most, but not all, the viable state space behavioral models (p<.05, Wilcoxon signed rank test; Fig. 2; **Supplemental Figure 7**).

When modeling Conflict versus Adaptation states, significant differences in stimulation-related effects on reaction time were found (Fig. 2; **Supplemental Figure 7; Supplemental Table 4**). Stimulation of the amygdala resulted in significantly higher emotion conflict state (ECS) values compared to non-stimulated blocks, translating to an overall slower ECS-driven RT component (*p=0.0371*; Wilcoxon signed rank test; Model 1), but no significant change in the RT component corresponded with the emotion adaptation state (EAS) (*p=0.9219*; Wilcoxon signed rank test; Model 1). In contrast, stimulation of the rACC resulted in a significantly faster reaction time EAS-component (*p=0.0005*; Wilcoxon signed rank test; Model 1). When modeling the equipoise between conflict (ECS) versus adaptation (EAS; Model 15), a state we labeled Conflict-Adaptation state (Fig. 2), stimulation of the dmPFC and rACC resulted in significantly positive deviations in Conflict-Adaptation equipoise, representing ECS-related slowing of RTs compared to the EAS-driven RTs (*p=0.0215* for dmPFC stimulation; *p=0.0025* for rACC stimulation; Wilcoxon signed rank test; Model 15). In contrast, stimulation of the dACC and rACC resulted in significant negative deviations in Conflict-Adaptation equipoise, representing relatively slower EAS-driven reaction times relative to ECS-driven reaction times (*p=0.0016* for dACC stimulation; Wilcoxon signed rank test; Model 15). The effects of dmPFC versus dACC stimulation were statistically separable upon *post hoc* testing **(Supplemental Figure 7;** p=0.0116; Kruskal-Wallis test; **Supplemental Table 4**).

### Adaptive stimulation to test bidirectional control

Based on these results, and to more directly test the hypothesis that these stimulation effects were bidirectional, we chose two regions, the dACC and dmPFC, to stimulate in a closed-loop, real-time setting. We chose the dmPFC over the rACC since stimulation in the rACC produced highly variable results on a per block level (Fig. 2F). Estimating the Conflict-Adaptation state on a trial-by-trial basis (Fig. 2D, 3), we alternately set the real-time algorithm to stimulate dmPFC when the Conflict-Adaptation was low, representing decreased RT to conflict relative to adaptation, then to stimulate the dACC when the state was high, representing increased RT to conflict relative to adaptation. In essence, this would create a “state clamp” which could maintain a state within specified bounds. In all three participants, dmPFC stimulation increased the Conflict-Adaptation state both during and after stimulated trials, representing an increase in RT to conflict trials relative to adaptation trials. In contrast, dACC stimulation decreased the Conflict-Adaptation state, representing a decrease in RT to conflict trials relative to adaptation trials (Fig. 3B-D). To examine the temporal extent of these changes, when we averaged the Conflict-Adaptation state for stimulated trials and the subsequent 4 trials, we found a significant difference between dmPFC and dACC stimulation in the Conflict-Adaptation state equipoise values in two of the three participants (EP23: p=0.0003 and EP24: p=0.0012; EP21: p=0.6438; Wilcoxon rank sum test; Fig. 3E-F). Two of the participants spontaneously volunteered their subjective impression of the adaptive stimulation testing following the final stimulation (Test) block. One participant (EP23) stated they felt like “the answers were more immediate…. [The image] shows up and it was a reactionary, I hit the button.” A second participant (EP24) stated “It was harder to, like, think… but the task seemed easier… I didn’t have to think about [the task] as much.”

**Figure 3.**
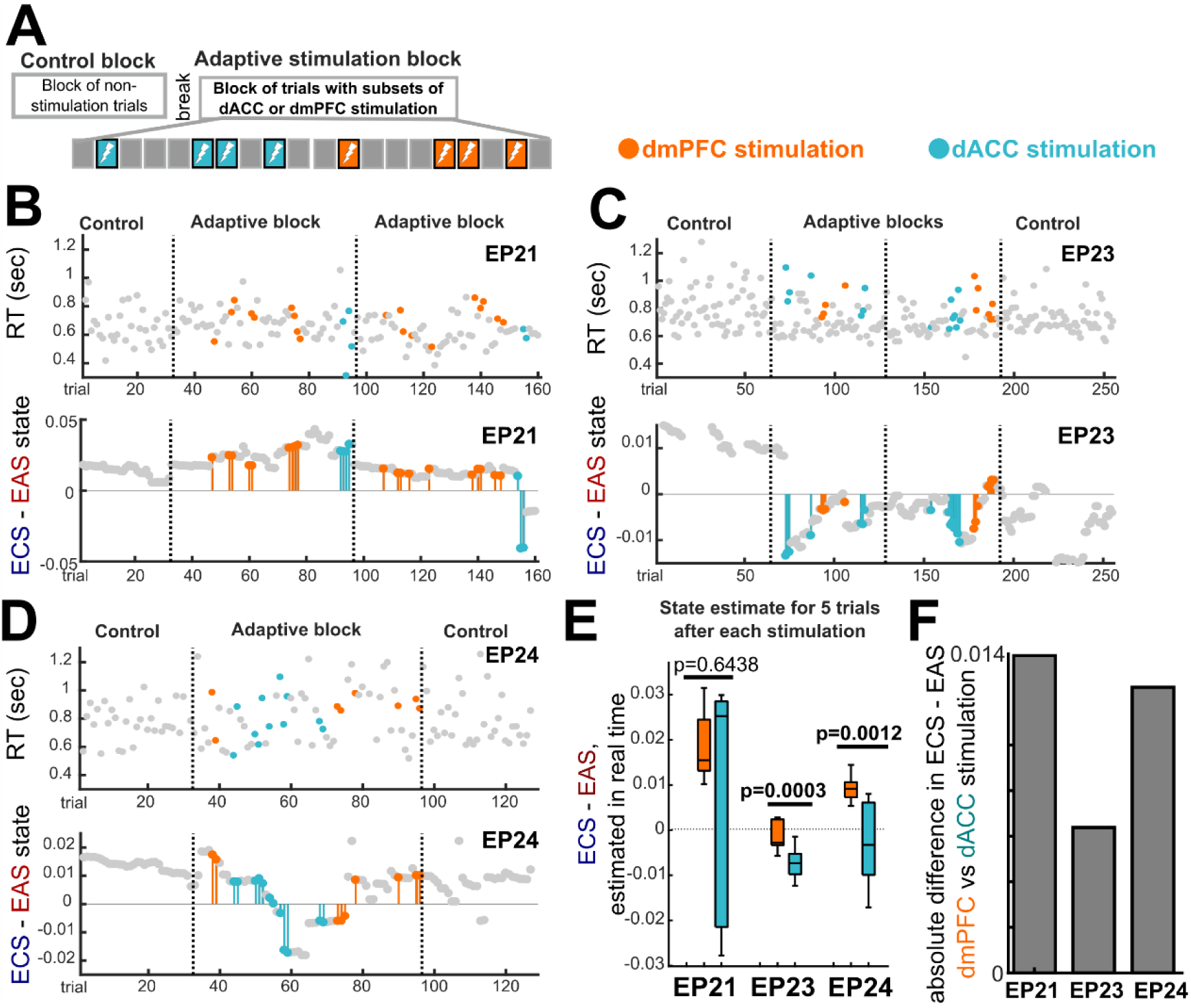
Real time estimate and bidirectional modulation of behavior using dmPFC and dACC stimulation. **A**. Experimental design. **B-D.** Top: Reaction time changes with dmPFC (orange) and dACC (teal) stimulated trials indicated by a color change (N=3). Bottom: The real-time Conflict-Adaptation state (ECS-EAS) during Control blocks (with no stimulation) and model-based Blocks with dmPFC (orange stems) and dACC (teal stems) stimulated trials. Grey dots: no stimulation. EPxx indicate the different participants. **E.** After averaging the current stimulated trial and the subsequent 4 trials per dmPFC stimulation (orange) versus per dACC stimulation (teal), we found a significant difference between dmPFC and dACC stimulation for EP23 (p=0.0003) and EP24 (p=0.0012), but not exceeding significance for EP21 (p=0.6438; Wilcoxon rank sum test). Error bars are standard error across trials. **F.** dmPFC stimulation induced higher Conflict-Adaptation state values when we subtracted the Conflict-Adaptation state with dACC versus dmPFC stimulated trials. Brain region abbreviations as in **Fig. 2**.

## DISCUSSION

The current study sought to model and modulate the hidden cognitive states associated with emotion regulation using a combination of state-space modeling of the behavior and intracranial stimulation. Using a behavioral state space approach and a well-validated emotion regulation task, we were able to isolate and model emotion conflict (ECS) and emotion adaptation states (EAS) with good model fit, convergence, and predictive ability. Using intracranial electric stimulation applied to emotion regulation-related neurocircuitry, we were able to bi-directionally modulate these state-dependent behavioral responses. Finally, we were able to use this state-space model to estimate changes in ECS-EAS equipoise (Conflict-Adaptation equipoise) and to modulate these states upwards or downwards using predictive, adaptive, closed-looped stimulation, with observable behavioral effects.

Taking a state-space approach to modeling behavior allowed for more precise mapping of relevant behavior to underlying cognitive processes. Using this approach with the ECR task, we were able to model and predict behavioral responses to emotion conflict (CI) and emotion conflict resolution (II) trials even during the presentation of alternative trial types (e.g. predicting how the participant would respond to a CI trial during a CC or IC trial). Importantly, the state estimate modeling approach allowed us to *predict* the Emotion Conflict (ECS) or the Emotion Adaptation (EAS) cognitive states. This represents a significant improvement over traditional behavioral modeling approaches, wherein behavioral responses to task conditions would be averaged by condition across the entire task, masking potentially important nuanced processes contributing to behavioral responses. Indeed, whereas our state-space approach was able to isolate specific features of ECR task trials, for example, trials that are directly affected by the immediately preceding trial (e.g. incongruent trials immediately preceded by a congruent trials (CI); incongruent trials immediately preceded by an incongruent trial (II)); taking a task condition-average approach did not reveal any significant differences in behavior between these trial types.

Having the ability to isolate and predict behavior based upon these hidden cognitive states associated with emotion regulation opens up the possibility for the development of novel neurotherapeutic approaches to modulate emotion regulation behavior. The features captured by the state-space approach in the current study - namely, the ability to detect emotion conflict and the ability to adapt behavior following conflict - have direct clinical implications and may be viable targets for intervention. For example, severe depression and anhedonia is associated with reduced reactivity to salient cues, potentially captured by deviations in the emotion conflict state (1, 2, 5, 48). By contrast, many disorders including ADHD, OCD, or GAD are marked by persistent perseveration and an inability to adapt and update behavioral responses (1, 2, 5, 34), potentially captured by deviations in the emotion adaptation state. Both these deviations from healthy, adaptive processing contribute to overall deficits in emotion regulation. Isolating and modulating these features of emotion regulation can provide a novel therapeutic approach in which these separable states are driven clamped in a more normal state in a patient-specific manner. Indeed, we have made progress recently in mapping behavioral states through this state estimate approach to neural activity in a related Stroop task (47) and mood can be decoded from neural data (35, 59), which means that we could, in theory, identify the neural signatures mapped to these underlying hidden states to then close the loop between stimulation and neural signatures of hidden cognitive states as surprise (ECS) or adaptation (EAS) (19–21).

We were able to demonstrate an ability to drive these states, and modulate behavior accordingly, using intracranial electric stimulation to a subset of brain regions implicated in emotion regulation, namely the dlPFC, dACC, rACC, dmPFC, and amygdala. This is a direct probe of causality between driving neural activity and the alteration of separate cognitive processes underlying behavior during the ECR task. Specifically, dlPFC stimulation resulted in significantly faster reaction times overall, whereas stimulation of the amygdala resulted in significantly slower reaction times overall. This finding is intriguing in light of the now well-established cortical control theory of emotion regulation, wherein increased dlPFC engagement is associated with decreased amygdala activation, representing cortical control over salience processing (26, 28–30). Location-specific effects were demonstrated both in the change in reaction times and behavioral changes within the state-space modeling framework. The state-space model demonstrated that stimulation in the dACC accelerated emotion conflict processing while stimulation in the dmPFC and rACC slowed emotion conflict processing. This was further demonstrated in a predictive fashion using closed loop stimulation in two of three participants.

The behavioral changes due to stimulation were illustrated within the context of the Conflict-Adaptation state, particularly for dmPFC and dACC stimulation. However, a key feature of our investigation is that we could independently modulate different hidden states indicating that not only are these states separate but that they are supported by, and can be influenced differently by, changes in activity in identified brain regions. For instance, dlPFC and rACC stimulation had effects on individual components of the state estimate model, with dlPFC stimulation decreasing reaction time bias. Interestingly, transcranial magnetic stimulation (TMS) of the dlPFC has been shown to not only modulate attention in a threat task, but this response varied with intrinsic anxiety levels in the participants (60). These results may correlate to our highly consistent effect of dlPFC direct electrical stimulation on reaction time bias (likely includes attentional components). It would be interesting if additional TMS approaches or other non-invasive neuromodulatory techniques could replicate our results from direct electrical stimulation, particularly with regard to stimulation in the dmPFC versus the dlPFC (61). Alternately, rACC stimulation altering the emotion conflict resolution (EAS) value of the state estimate model which was not surprising considering this area has been shown to be a key part of the brain network supporting emotion processing and regulation in general (6, 22, 26–28, 31) and stimulation in this region has alleviated depression in a subset of patients (62–64). Stimulation of the amygdala resulted in significantly slower reaction times during emotion conflict, but not during emotion conflict resolution, suggesting increased processing of emotion salience during emotion conflict. Therefore, we were able to isolate and modulate different features of the behavior, such as attention or the effects of trial history, using the state estimate approach. In addition, the state space framework allowed us to address other features of the behavior, such as unexpected changes in trial types (e.g. Congruent to Incongruent trials). The Conflict-Adaptation model of behavior revealed a bidirectional effect of dmPFC/rACC versus dACC stimulation, both on the full block level and when implemented in real time in closed-loop control (dmPFC and dACC). Thus, the current study was able to isolate clinically-relevant behavior and was able to modulate this behavior in real time.

### Limitations

There are several limitations to the interpretation of the current study. First, whereas we were able to demonstrate the ability to model and modulate behavior in an emotion regulation task, the ecological validity of this approach remains to be tested. Future studies are needed to examine whether the benefits of stimulation parameters modeled based upon a behavioral task can be extended to functioning in the real world. Second, the current study relied upon a sample of convenience (patients undergoing clinical monitoring for epileptic seizures). Thus, electrode placement was limited to the clinical specifications of this sample. There may be other, more effective targets for modulating emotion regulation, such as the ventrolateral prefrontal cortex or inferior parietal lobule (26, 27, 29, 31). Future studies are needed to identify optimal targets for stimulation. Third, in order to implement the state space behavioral approach of characterizing hidden cognitive states in a clinical setting, we would need to account for the wide variance in both the stimulation induced changes in reaction times and, to a lesser extent, in the state estimate values. We hypothesized this variance could be due to the noisy signal inherent to reaction time (40, 47), the temporal rarity of the sampling of state (we only captured behavior in discrete trials over time), and variability in electrode positions, as they were placed for clinical purposes related to epilepsy and not to specifically target subregions of cingulate, prefrontal cortex or other areas related to emotional control. Future studies are needed to disentangle these potential sources of variance.

### Conclusion

The current study represents a significant step towards delineating and modulating identified and dissociable, relevant hidden cognitive features associated with emotion regulation using intracranial neuromodulation at multiple brain regions. Further, it is, to our knowledge, the first study of its kind to demonstrate closed-loop modulation of emotion regulation. These results suggest a potential pathway towards the development of novel neurotherapeutics by leveraging our understanding of neural activity from fMRI from previous studies with the use of neural stimulation and multiple metrics of behavior (psychometric questionnaires, behavioral tasks, and state estimate modeling) to arrive at a therapeutic strategy, in this instance for emotion dysregulation. In this context, the possibility bidirectional altering of emotion regulation-related behavior with dmPFC/rACC versus dACC as outlined in the current study suggests a therapeutic approach that can help maintain an optimal balance of emotion processing and prevent behavior from being driven too far toward either end of the continuum. These results, therefore, could provide an informed basis for the use of intermittent and targeted neuromodulation to aid individuals experiencing severe emotion dysregulation at both extremes of the spectrum. Indeed, this approach, applied to other domains and across cognitive and emotional tasks, could allow us to arrive at a more refined view of how to use neural stimulation to therapeutically alter the circuitry underlying important domains of functioning, such as maladaptive emotional processing and decision making, with implications for a wide array of neuropsychiatric diseases (33, 34, 51, 52).

## Acknowledgments

We thank Giovanni Piantoni, Jean-Baptiste Eichenlaub, Pariya Salami, Erica Johnson, Chris Salthouse, and Mia Borzello for their help in acquiring and analyzing the data.

## Author contributions

Conceptualization, A.C.P., U.T.E., A.S.W., T.D., D.D.D, K.K.E., E.N.E., S.S.C.; Methodology: A.C.P., S.S.C., A.S.W., N.N., K.E.E., B.C., S.Z., A.Y., U.T.E., T.D., D.D.D., E.N.E.; Resources, S.S.C., D.D.D., E.N.E.; Investigation, K.F., A.C.P., B.C., D.V.-L., G.B., S.Z., A.A., A.G., K.K.E., A.S.W., S.S.C.; Formal Analysis, A.C.P.; Visualization, A.C.P.; Writing-Original Draft, A.C.P., S.S.C.; Writing-Review & Editing, A.C.P., S.S.C., A.S.W., A.Y., K.F., K.K.E., G.B., A.E., D.D.D., E.N.E.; Funding Acquisition, A.S.W., T.D., D.D.D, E.N.E., S.S.C.; Supervision, U.T.E., S.S.C.

## Supplemental Material

Supplemental Figures 1-7

Supplemental Tables 1-4

## Materials and Methods

### Participants

All participants provided fully informed consent according to NIH and Army HRPO guidelines as monitored by the local MGH Institutional Review Board. ECR task data from three separate cohorts (N=99) were included in the state-space models: healthy control volunteers (*n*=42, mean age = 32.9, range = 20-55, 36% female); psychiatric patients (*n*=16, mean age = 33, range = 20-59, 62.5% female; see **Supplemental Table 1** for diagnostic details), and patients with epilepsy (*n*=41, mean age = 36.15, range = 14-68, 65% female). Only the patients with long-standing pharmaco-resistant complex partial seizures were included in the subsequent intracranial stimulation study in the context of ongoing clinical care, and three of these patients participated in the closed-loop (adaptive) stimulation trial. The patients with epilepsy were implanted with electrodes as part of a course of clinical monitoring using intracranial electroencephalogram (iEEG) recordings. Patients with epilepsy were implanted with multi-lead depth or grid electrodes (a.k.a iEEG) to confirm the hypothesized seizure focus, and locate epileptogenic tissue in relation to essential cortex, thus directing surgical treatment (**Supplemental Table 3**). The decision to implant electrodes, the number, types and location of the implantations were all determined on clinical grounds by a team of caregivers independent of this study. Participants with epilepsy were informed that participation in the current study would not alter their clinical treatment in any way, and that they may withdraw at any time without jeopardizing their clinical care. Healthy control participants and participants diagnosed with psychiatric disorders completed the behavioral task but were not implanted with any electrodes. All participants voluntarily participated after fully informed consent according to NIH and Army HRPO guidelines as monitored by the local MGH Institutional Review Board.

### Behavioral Task

The ECR task is a well-validated task designed to assess the effects of emotional conflict that arises from the incompatibility between task-relevant and task-irrelevant emotional dimensions of a stimulus. Faces with fearful and happy expressions are presented with the words “happy” or “fear” written across them. Words are either *congruent* (e.g. “happy” written across an image with a happy expression) or *incongruent* (e.g. “happy” written across an image with a fearful expression; Fig. 1A; Subjects are asked to identify the emotional expression of the face while ignoring the word. Thus, successful completion of the task requires regulation of responses to task irrelevant emotional stimuli in order to focus on task relevant goals. Trials can be analyzed with regard to immediately preceding trials: incongruent trials preceded by congruent trials (CI trials) measure emotion conflict, and incongruent trials preceded by incongruent trials (II trials) measure resolution of emotion conflict. The images were presented in a pseudorandom order such that the identity, gender, and valence were shown randomly. Congruence changes were balanced in that there were no more than three congruent or incongruent trials in sequential order. Task stimuli were presented with either Presentation software (Neurobehavioral Systems) or Psychophysics toolbox in MATLAB (65–67) (Mathworks, Natick, MA). The task in the epilepsy monitoring unit (EMU) was composed of at least one and up to six 64-trial blocks of images presented for 1 second, with a fixation cross presented for 2-4 seconds in between images. The task performed outside of the EMU included one 152 trial block collected during MEG/EEG monitoring (data outside the scope of the current study). Aside from the number of trials, the timing of the task trials was the same between settings. Behavioral data were analyzed using MATLAB (Mathworks, Natick, MA) and consisted of reaction times and response accuracy.

### Psychometric Questionnaires

In addition to the ECR behavioral task, all participants were asked to complete a set of self-report questionnaires assessing emotion reactivity and regulation ability. They completed Emotional Reactivity Scale (ERS) is a 21-item self-report measure designed to address emotion sensitivity, intensity and persistence (55). The Difficulties in Emotion Regulation Scale (DERS) is a 36-item self-report measure reflecting difficulties in emotional understanding and awareness as well as the acceptance of emotions and the ability to refrain from impulsive behavior when experiencing negative emotions (8). The Anxiety Sensitivity Index (ASI) is a 16-item questionnaire that measures an individual’s concern about the possible negative consequences of anxiety symptoms (56, 57, 68). We used the average and standard deviation of the 42 healthy control participant score response to zscore all the participant responses using the distribution of healthy control participant responses as the reference group. This was done to allow for the comparison and combination of substantially different score values from the psychometric questionnaires and to correlate values with the behavioral data.

### Developing and Validating Models of ECR Hidden States

Using only the subset of data from sessions of ECR without any direct electrical stimulation and the COMPASS state-space toolbox ((40); https://github.com/Eden-Kramer-Lab/COMPASS), we derived a set of behavioral state-space models of underlying hidden states for ECR. The model structure was similar between models with differences being what features of the task and responses from the participants were used to generate the model. Choices in the model parameters and features (e.g. trial type) were informed by the existing ECR literature to reflect emotion conflict and emotion conflict resolution (6, 19–23, 48). Included features of the task were whether we include the reaction time, reaction times during specific trial types (such as incongruent vs congruent trials), and accuracy (correct/incorrect; **Supplemental Figure 3**).

To illustrate how we built each state estimate model, we will demonstrate how we modeled Conflict-Adaptation (Fig. 1E; **Supplemental Figure 4**) informed by the current thinking that adaptation to Incongruent-Incongruent (II) trial sequences was captured in an emotion adaptation state (EAS), whereas the switch burden of Congruent-Incongruent (CI) trial sequences was captured as an emotion conflict state (ECS; (6, 23)). The model was composed of three main components which varied trial to trial: trial-independent, baseline reaction time or drift state, also called the RTB (X_*k,bias*_), ECS (CI trials; X_*k,CI*_) and EAS (II trials; X_*k,II*_). We used these three state estimates to determine both overall and trial-to-trial changes in reaction time behavior, separating the effects of drifts in attention or distraction (the bias term RTB) from the effects of the relevant trial types (CI or II) on behavior, which we could observe in the resulting state estimates on a trial by trial basis (Fig. 1 D). The difference between ECS and EAS would then reflect the balance of the effects of surprise related to a change in conflict on behavior versus adapting to conflict. Therefore, we could regress out latent variables likely corresponding to hidden cognitive states as well as identify interactions between these independent latent variables (**Supplemental Figure 3-4**).

Since bias, CI, and II trials do not fully describe all features of ECR, to more fully explore the hidden features which could be underlying ECR task behavior, we ran 30 additional models on the behavioral data set with different state parameters using the state estimate approach using the following metrics to compare the models: 1) Interpretability: whether the terms in the model could be mapped to independent features of the task or behavior; 2) Convergence: if the models could converge using expectation maximization (EM) and maximum (ML) likelihood measures; 3) Variability: explanation of the variability without over-specifying the number of terms needed to predict behavior (relevant to the noise terms); 4) Predictability: identifying which models best predicted reaction time; and 5) Psychometric correspondence: correlation of an underlying emotion reactivity state with self-report questionnaires (**Supplemental Figure 2-4**).

The goal of this work was to arrive at one or a subset of validated state estimate models which quantify changes in hidden cognitive states, and which operate across a population of participants from the non-stimulated ECR behavioral task sessions (N=99). Therefore, each of these measurements were examined in turn using the data set from 99 individuals performing the ECR task both in and outside the EMU, exclusively with data from task sessions without neural stimulation (**Supplemental Figure 3-4**). We also confirmed the models could converge for both larger and smaller data sets with tolerable noise ranges (data not shown).

Model interpretability directly related to chosen model features informed by both the behavior during the task and the literature surrounding Stroop and ECR tasks (**Supplemental Figure 3-4;** (6, 19–22)). For instance, several models have a term for reaction time bias, a trial-independent, baseline reaction time or drift state, which could correspond to an overall attention signal or effort. Other models include trial feature-relevant changes in behavior. In addition, some models include a state variable measuring how reaction time is influenced by the transition term (Tr.), namely a term indicating whether the previous trial was different to the current trial versus the two trials being the same (**Supplemental Figure 3-4**), a correlate of cognitive flexibility from trial to trial. Goal-oriented and conflict resolution hidden states could be addressed by a state variable related to whether the current trial is congruent or incongruent (CvsI), which details the effects of the current trial difficulty on reaction time which is represented in several models as well (**Supplemental Figure 3-4**). Finally, we included models which incorporate state variables related to the valence of the face or the word (**Supplemental Figure 3-4**). In addition to the trial types, state estimate models could also incorporate and predict different behavioral components such as reaction time or accuracy (**Supplemental Figure 3-4**). We developed reaction time-only behavioral models, accuracy-only models, and mixed models (**Supplemental Figure 3-4**).

A key point to creating these models is that we had to use indicator (e.g. trial type) and state estimate variables which could be independent of one another. For instance, we could not include indicator terms which were mutually exclusive, such as whether the trial was congruent or incongruent, as separate state estimate terms. Second, we also tested whether we could ‘fix’ some model terms such that, in the process of estimating the coefficients for each model to describe the behavior, we did not iteratively determine the state per trial. For instance, when we identified the congruence term as ‘fixed’, this means that we assumed the behavioral responses to trial congruence does not vary from trial to trial. The fixed terms as well as the per trial state space variability were both tested along with the different types of trial features to determine if we could identify both the least number of features and what features in a state space model could be used to describe and predict behavior. Of course, considering the large number of different parameters we could use to model behavior (**Supplemental Figure 3-4**), we could be overfitting the behavior with too many interrelated and dependent terms. For this reason, we generated 30 separate models so that, in each model, the state variable terms, behavioral measure, and trial-relevant features used in the models could be independent from one another in each formulation while minimizing the number of state estimates used per model (**Supplemental Figure 3-4**).

#### Convergence Varies Between Models

Following feature selection to generate these state estimate models, we then performed an expectation maximization (EM) algorithm to measure the maximum likelihood estimates of the model parameters and filter solutions per task run to arrive at task run-specified parameters. We addressed convergence by identifying whether the models converged on a solution as measured by their maximum likelihood estimate. We found many of the state space models converged on a solution, though some did not (**Supplemental Figure 3-5**). Setting a criterion threshold to reject models, we could eliminate six state space models since they took more iterations to converge or never converged on a stable solution, as reflected by the lower maximum likelihood slopes in the first 500 iterations after normalizing to the maximum per curve (p<0.000001; Chi-sq= 1,053; Friedman test; **Supplemental Figure 3-5**). Most of the models that failed to converge involved fixing the reaction time bias (or reaction time drift) to a constant value across trials or estimating reaction time bias without taking into account trial transitions or trial types (**Supplemental Figure 3-5**).

#### Model Variability and Noise Can Used to Reject Models

The models were compared based on the width of the confidence bounds of each state estimate, a variability term W_k_ (extent of the state process noise per each state variable), and an observed behavior (RT) noise V (for the estimated noise of the model, **Supplemental Figure 3-5**). We identified which models had the least predicted observation noise and per state variable (**Supplemental Figure 4**). Model 5 had no estimate of the reaction time bias state and the predicted observation noise was significantly higher than for other models (V; p<0.000001; Chi-sq= 1,225; Friedman test; **Supplemental Figure 4**). For each state variable, per-state estimate noise (W_k_) was higher for an additional five models (Models 7, 9, 11, 20, and 14), thereby eliminating them from the list (p<0.000001; Chi-sq= 20,663; Friedman test; **Supplemental Figure 4**). These rejected models, interestingly, did not contain an adaptation (II) term.

#### Models Vary in the Behavioral Prediction

As the models could be used to predict reaction times per each trial (40, 41), we performed a leave-one-out cross validation test where we predicted behavior (reaction time) by iteratively removing one trial, performing the model fit to predict that reaction time, and then moving on to the next trial. We then calculated the root mean square (RMS) difference between the actual and predicted reaction times and compared results across models. This step allowed us to determine how well each state space model could predict behavior. To test the predictive power of each model, we ran the model fitting iteratively, removing a single trial but including all the other trials iteratively across all trials and using the censor capabilities of the COMPASS package to replace, or predict, the missing reaction time per session across the data set (40). We found that most of the models could predict reaction times at similar levels. In fact, the differences between the actual and predicted log reaction times were not significantly different between models except for the significantly lower differences for Models 6 and 8 and significantly higher values for Model 5 (p<0.00001; Friedman test; **Supplemental Figure 4**), resulting in model rejection.

#### Models Using Reaction Time Can Describe the Behavior Without Accuracy

While we did include models which had binomial state variables which took into account trial accuracy (**Supplemental Figure 3**), we found that models which only included trial accuracy (Models 23-26) either did not converge or had very high confidence bounds and noise terms. Models which included both reaction time and trial accuracy, interestingly, were not different in the state estimates to when we included only the reaction time in the models (data not shown). We took this result to mean that the high accuracy (88% correct) and the intermittent incorrect trials (Fig. 1) were not informative to drive these hidden cognitive states. For this reason, we did not include these models (Models 23-30) in the remaining results.

#### State Variables Correspond with Psychometric Questionnaire scores

We also tested whether these state estimates correlated with psychometric questionnaire scores (**Supplemental Fig. 2, 4**). We calculated the correlation between the state space variables per model and each psychometric scale z-scored relative to the healthy control group using only the data set when there was no stimulation in the entire session (**Supplemental Figure 4**).

### Intracranial (open-loop) stimulation

As part of clinical monitoring for seizures, patients with epilepsy were implanted with depth electrodes (Ad-tech Medical, Racine WI, USA, or PMT, Chanhassen, MN, USA) with diameter ranges of 0.8–1.0 mm and consisted of 8-16 platinum/iridium-contact leads at between 1-2.4 mm long. Electrodes were localized by using a volumetric image coregistration procedure. Using Freesurfer scripts (69, 70) (http://surfer.nmr.mgh.harvard.edu), the preoperative T1-weighted MRI (showing the brain anatomy) was aligned with a postoperative CT (showing electrode locations). Electrode coordinates were manually determined from the CT and placed into the native space (71). Mapping to brain regions was performed using an electrode labeling algorithm (ELA; (72); https://github.com/pelednoam/ieil). We mapped electrodes to regions in a given location which can be flexibly chosen within Freesurfer, using the DKT atlas in combination with a subcortical mapping (69, 70, 73).

To alter behavior in the ECR task using open loop stimulation, we used brief trains of focal high frequency open loop electrical stimulation (50, 51, 74) targeting the dACC, rACC, amygdala, dmPFC, and dlPFC in 16 of the total 41 participants with epilepsy where stimulation was performed during single, alternating blocks per brain region. Specifically, participants performed 64-trial blocks (Test Blocks) during which 130 Hz (N=2) or 160 Hz (N=14) stimulation for 400 ms was delivered at image onset at 4-6 mA at a single site. Stimulation was delivered in a bipolar configuration using CereStim (Blackrock, Salt Lake City, Utah). Pulses were charge balanced with a 90 µsecond negative deflection, a 53 µsecond interval and a 90 µsecond positive deflection. During Test Blocks, stimulation trials were interspersed with non-stimulation trials (Fig. 2A, **Supplemental Table 1; Supplemental Figure 6**). In the same session, participants also performed 64-trial blocks without stimulation (Control Blocks, Fig. 2A).

In all cases, we performed safety tests of multiple levels of current in each stimulation site to look for signs of epileptogenic activity as well as to ask participants if they experienced any sensations with stimulation. Only if there was neither epileptiform activity nor subjective awareness of the stimulation did we perform the task. We did not include a forced choice task to determine that there was no subjective effect. We balanced the ECR stimulation protocol to study within-testing effects of neural stimulation on the reaction times of the participants in alternating blocks and trials. labeled. Each participant was unaware if the stimulation was occurring in a particular trial or block. To examine changes in reaction time or state estimates due to stimulation, we z-scored all reaction times relative to the Control blocks to be able to compare across participants.

### Adaptive (closed-loop) stimulation

We used our Model 1, which we call the Conflict-Adaptation state (Fig. 1), to track the Conflict-Adaptation equipoise in real time and to deliver during adaptive stimulation in two separate sites, the dmPFC and the dACC. The dmPFC and the dACC were chosen based on the open loop stimulation results. These two sites were selected based upon the results of our intracranial stimulation experiments (see Results section). The computer displaying the ECR task was interfaced via the Psychophysics toolbox with a separate computer. The interface registered the trial type, trial history, and the reaction time on a trial by trial basis as the participants performed the task, using a National Instruments digital data acquisition board (NI USB-6501; NI) and a custom MATLAB program to calculate the Conflict-Adaptation state in real time. This closed-loop approach involved ‘training’ the model on prior non-stim ECR task sessions from the same participant to arrive at a convergent estimate of model parameters with 1000 iterations. We validated this approach by applying the same model estimates for sessions across multiple days and found generally similar state estimates could be arrived at per patient across days (data not shown). We chose 1000 iterations since this was well beyond the point when the maximum likelihood value stabilized, or reached an asymptote, following the expectations maximization step while the maximum likelihood did not always asymptote at 500 iterations (**Supplemental Figure 4-5**). The resultant model and model parameters could then be used in real time such that, for each trial, the resultant emotion conflict state (ECS), emotion adaptation state (EAS), and reaction time bias (RTB) values could be estimated. We hypothesized that we could use dmPFC and dACC stimulation to bidirectionally alternate the Conflict-Adaptation (ECS>EAS) state based on the opposite effects of dACC and dmPFC stimulation during open loop stimulation (Fig. 2). To test this, we applied stimulation based on the real-time values for the Conflict-Adaptation state. We alternately set the real-time algorithm to stimulate dmPFC when the Conflict-Adaptation state was below zero to drive the value upward closer to zero, then to stimulate dACC while the state was above zero to drive the value downward. In essence, this would create a “state clamp” which could maintain a state within specified bounds, with zero representing equipoise between the conflict and adaptation state. The primary goal was to alternate between stimulation sites to determine if we could shift the Conflict-Adaptation state bidirectionally within the same task block. Further analyses of the behavior involved taking the dACC and dmPFC stimulation trials and the 4 trials subsequent to the stimulation, averaging these four sequential trials, and comparing dynamics between the dACC and dmPFC stimulation trials.

### Statistical analysis

All statistical comparisons were performed using non-parametric statistical approaches. We tested comparisons across brain regions with the Kruskal–Wallis test for non-equivalence of multiple medians followed by *post hoc* Tukey-Kramer tests to determine statistically separable groups. For correlations to reaction time or changes in z-scored reaction times relative to Control blocks, we used a Wilcoxon signed rank test within brain regions to determine if a distribution’s mean was significantly different from zero. For comparisons between Control and Test blocks in power and reaction time, we used the Wilcoxon rank sum test (two-sided) for comparisons between individual medians. We additionally used the Wilcoxon signed rank test (two-sided) for determining if a distribution’s median was significantly different from zero.

### Data and materials availability

All data underlying the study is available upon request.

### Code availability

All code, including custom MATLAB code, underlying the study will be made available upon request.

## References

1. J. J. Gross, The emerging field of emotion regulation: An integrative review. Rev. Gen. Psychol. 2, 271–299 (1998).

2. L. M. Braunstein, J. J. Gross, K. N. Ochsner, Explicit and implicit emotion regulation: A multi-level framework. Soc. Cogn. Affect. Neurosci. 12, 1545–1557 (2017).

3. E. Halperin, Emotion, Emotion Regulation, and Conflict Resolution. Emot. Rev. 6, 68–76 (2014).

4. S. L. Koole, The psychology of emotion regulation: An integrative review. Cogn. Emot. 23, 4–41 (2009).

5. J. J. Gross, Handbook of Emotion Regulation (Guilford Press, 2014).

6. A. Etkin, T. Egner, D. M. Peraza, E. R. Kandel, J. Hirsch, Resolving emotional conflict: a role for the rostral anterior cingulate cortex in modulating activity in the amygdala. Neuron 51, 871–82 (2006).

7. K. N. Ochsner, J. A. Silvers, J. T. Buhle, Functional imaging studies of emotion regulation: A synthetic review and evolving model of the cognitive control of emotion. Ann. N. Y. Acad. Sci. 1251, E1–E24 (2012).

8. K. L. Gratz, L. Roemer, Multidimensional Assessment of Emotion Regulation and Dysregulation: Development, Factor Structure, and Initial Validation of the Difficulties in Emotion Regulation Scale. J. Psychopathol. Behav. Assess. 26, 41–54 (2004).

9. J. C. Fowler, et al., Emotion dysregulation as a cross-cutting target for inpatient psychiatric intervention. J. Affect. Disord. 206, 224–231 (2016).

10. E. Halperin, Emotion, Emotion Regulation, and Conflict Resolution. Emot. Rev. 6, 68–76 (2014).

11. M. Berking, et al., Deficits in emotion-regulation skills predict alcohol use during and after cognitive behavioral therapy for alcohol dependence. J. Consult. Clin. Psychol. 79, 307–318 (2011).

12. X. Caseras, et al., Emotion regulation deficits in euthymic bipolar I versus bipolar II disorder: a functional and diffusion-tensor imaging study. Bipolar Disord. 17, 461–470 (2015).

13. R. A. Lanius, et al., Emotion modulation in PTSD: clinical and neurobiological evidence for a dissociative subtype. Am. J. Psychiatry 167, 640–647 (2010).

14. D. S. Mennin, R. M. Holaway, D. M. Fresco, M. T. Moore, R. G. Heimberg, Delineating Components of Emotion and its Dysregulation in Anxiety and Mood Psychopathology. Behav. Ther. 38, 284–302 (2007).

15. J. Townsend, L. L. Altshuler, Emotion processing and regulation in bipolar disorder : a review. Bipolar Disord. 14, 326–339 (2012).

16. M. T. Tull, H. M. Barrett, E. S. Mcmillan, L. Roemer, A preliminary investigation of the relationship between emotion regulation difficulties and posttraumatic stress symptoms. Behav. Ther. 38, 303–313 (2007).

17. N. H. Weiss, M. T. Tull, M. D. Anestis, K. L. Gratz, The relative and unique contributions of emotion dysregulation and impulsivity to posttraumatic stress disorder among substance dependent inpatients. Drug Alcohol Depend. 128, 45–51 (2014).

18. C. E. Wilcox, J. M. Pommy, B. Adinoff, Neural circuitry of impaired emotion regulation in substance use disorders. Am. J. Psychiatry 173, 344–361 (2016).

19. T. Egner, Congruency sequence effects and cognitive control. Cogn. Affect. Behav. Neurosci. 7, 380–390 (2007).

20. T. Egner, Surprise! A unifying model of dorsal anterior cingulate function? Nat. Neurosci. 14, 1219–1220 (2011).

21. T. Egner, A. Etkin, S. Gale, J. Hirsch, Dissociable neural systems resolve conflict from emotional versus nonemotional distracters. Cereb. Cortex 18, 1475–84 (2008).

22. A. Etkin, T. Egner, R. Kalisch, Emotional processing in anterior cingulate and medial prefrontal cortex. Trends Cogn. Sci. 15, 85–93 (2011).

23. A. Etkin, C. Buechel, J. J. Gross, The neural bases of emotion regulation. Nat. Rev. Neurosci. 16, 693–700 (2015).

24. A. Etkin, Neurobiology of anxiety: from neural circuits to novel solutions? Depress. Anxiety 29, 355–358 (2012).

25. A. S. Heller, et al., Reduced capacity to sustain positive emotion in major depression reflects diminished maintenance of fronto-striatal brain activation. Proc. Natl. Acad. Sci. 106, 22445–22450 (2009).

26. M. Comte, et al., Dissociating Bottom-Up and Top-Down Mechanisms in the Cortico-Limbic System during Emotion Processing. Cereb. Cortex 26, 144–155 (2016).

27. A. S. Eden, et al., Emotion regulation and trait anxiety are predicted by the microstructure of fibers between amygdala and prefrontal cortex. J. Neurosci. 35, 6020–6027 (2015).

28. S. J. Banks, K. T. Eddy, M. Angstadt, P. J. Nathan, K. Luan Phan, Amygdala-frontal connectivity during emotion regulation. Soc. Cogn. Affect. Neurosci. 2, 303–312 (2007).

29. F. Chen, et al., Increased Inhibition of the Amygdala by the mPFC may Reflect a Resilience Factor in Post-traumatic Stress Disorder: A Resting-State fMRI Granger Causality Analysis. Front. Psychiatry 9, 1–12 (2018).

30. W. W. Seeley, et al., Dissociable intrinsic connectivity networks for salience processing and executive control. J. Neurosci. 27, 2349–2356 (2007).

31. C. Morawetz, S. Bode, J. Baudewig, H. R. Heekeren, Effective amygdala-prefrontal connectivity predicts individual differences in successful emotion regulation. Soc. Cogn. Affect. Neurosci. 12, 569–585 (2017).

32. A. W. Laxton, N. Lipsman, A. M. Lozano, Deep brain stimulation for cognitive disorders, 1st Ed. (Elsevier B.V., 2013).

33. A. M. Lozano, N. Lipsman, Probing and Regulating Dysfunctional Circuits Using Deep Brain Stimulation. Neuron 77, 406–424 (2013).

34. A. S. Widge, et al., Treating refractory mental illness with closed-loop brain stimulation: Progress towards a patient-specific transdiagnostic approach. Exp. Neurol. 287, 461–472 (2017).

35. V. R. Rao, et al., Direct Electrical Stimulation of Lateral Orbitofrontal Cortex Acutely Improves Mood in Individuals with Symptoms of Depression. Curr. Biol. 28, 3893–3902.e4 (2018).

36. A. S. Widge, D. D. Dougherty, Deep Brain Stimulation for Treatment-Refractory Mood and Obsessive-Compulsive Disorders. Curr. Behav. Neurosci. Rep. 2, 187–197 (2015).

37. A. S. Widge, C. T. Moritz, “Closed-loop stimulation in emotional circuits for neuro-psychiatric disorders” in (Elsevier, 2016).

38. A. S. Widge, D. A. Malone, D. D. Dougherty, Closing the Loop on Deep Brain Stimulation for Treatment-Resistant Depression. Front. Neurosci. 12, 175 (2018).

39. S. Yamawaki, G. Okada, Y. Okamoto, I. Liberzon, Mood dysregulation and stabilization: Perspectives from emotional cognitive neuroscience. Int. J. Neuropsychopharmacol. 15, 681–694 (2012).

40. A. Yousefi, et al., COMPASS : An Open-Source, General-Purpose Software Toolkit for Computational Psychiatry. 12 (2019).

41. A. Yousefi, et al., Cognitive state prediction using an EM algorithm applied to Gamma distributed data. Conf. Proc. Annu. Int. Conf. IEEE Eng. Med. Biol. Soc. IEEE Eng. Med. Biol. Soc. Annu. Conf. 2015, 7819–7824 (2015).

42. J. Taghia, et al., Uncovering hidden brain state dynamics that regulate performance and decision-making during cognition. Nat. Commun. 9, 1–19 (2018).

43. M. J. Prerau, et al., Characterizing learning by simultaneous analysis of continuous and binary measures of performance. J. Neurophysiol. 102, 3060–3072 (2009).

44. M. J. Prerau, et al., A mixed filter algorithm for cognitive state estimation from simultaneously recorded continuous and binary measures of performance. Biol. Cybern. 99, 1–14 (2008).

45. A. C. Smith, S. Wirth, W. a Suzuki, E. N. Brown, Bayesian analysis of interleaved learning and response bias in behavioral experiments. J. Neurophysiol. 97, 2516–2524 (2007).

46. C. Martinez-Rubio, A. C. Paulk, E. J. McDonald, A. S. Widge, E. N. Eskandar, Multimodal Encoding of Novelty, Reward, and Learning in the Primate Nucleus Basalis of Meynert. J. Neurosci. 38, 1942 (2018).

47. A. Yousefi, et al., Decoding Hidden Cognitive States From Behavior and Physiology Using a Bayesian Approach. Neural Comput. 31, 1751–1788 (2019).

48. G. A. Fonzo, et al., Brain regulation of emotional conflict predicts antidepressant treatment response for depression. Nat. Hum. Behav. (2019) https:/doi.org/10.1038/s41562-019-0732-1.

49. J. R. Stroop, Studies of interference in serial verbal reactions. J. Exp. Psychol. 18, 643–662 (1935).

50. S. K. Bick, et al., Caudate stimulation enhances learning. Brain, 2930–2937 (2019).

51. Y. Ezzyat, et al., Closed-loop stimulation of temporal cortex rescues functional networks and improves memory. Nat. Commun. 9 (2018).

52. M. M. Shanechi, Brain – machine interfaces from motor to mood. Nat. Neurosci. 22, 1554–1564 (2019).

53. B. D. Greenberg, et al., Deep brain stimulation of the ventral internal capsule/ventral striatum for obsessive-compulsive disorder: worldwide experience. Mol. Psychiatry 15, 64–79 (2010).

54. B. D. Greenberg, et al., Three-Year Outcomes in Deep Brain Stimulation for Highly Resistant Obsessive–Compulsive Disorder. Neuropsychopharmacology 31, 2384–2393 (2006).

55. M. K. Nock, M. M. Wedig, E. B. Holmberg, J. M. Hooley, The Emotion Reactivity Scale: Development, Evaluation, and Relation to Self-Injurious Thoughts and Behaviors. Behav. Ther. 39, 107–116 (2008).

56. R. A. Peterson, S. Reiss, Test manual for the anxiety sensitivity index (International Diagnostic Systems, 1987).

57. B. J. Deacon, J. S. Abramowitz, C. M. Woods, D. F. Tolin, The Anxiety Sensitivity Index - Revised: Psychometric properties and factor structure in two nonclinical samples. Behav. Res. Ther. 41, 1427–1449 (2003).

58. J. Deng, G. Luan, Mechanisms of Deep Brain Stimulation for Epilepsy and Associated Comorbidities. Neuropsychiatry S(1), 31–37 (2017).

59. O. G. Sani, et al., Mood variations decoded from multi-site intracranial human brain activity. Nat. Biotechnol. 36, 954 (2018).

60. L. Sagliano, F. D’Olimpio, F. Panico, S. Gagliardi, L. Trojano, The role of the dorsolateral prefrontal cortex in early threat processing: A TMS study. Soc. Cogn. Affect. Neurosci. 11, 1992–1998 (2016).

61. J. A. Camprodon, A. Pascual-Leone, Multimodal Applications of Transcranial Magnetic Stimulation for Circuit-Based Psychiatry. JAMA Psychiatry 73, 407–408 (2015).

62. H. S. Mayberg, et al., Deep brain stimulation for treatment-resistant depression. Neuron 45, 651–660 (2005).

63. A. M. Lozano, et al., A multicenter pilot study of subcallosal cingulate area deep brain stimulation for treatment-resistant depression. J. Neurosurg. 116, 315–322 (2011).

64. A. Veerakumar, et al., Field potential 1/f activity in the subcallosal cingulate region as a candidate signal for monitoring deep brain stimulation for treatment-resistant depression. J. Neurophysiol. 122, 1023–1035 (2019).

65. D. H. Brainard, The Psychophysics Toolbox. Spat. Vis. 10, 433–436 (1997).

66. D. G. Pelli, The VideoToolbox software for visual psychophysics: Transforming numbers into movies. Spat. Vis. 10, 437–442 (1997).

67. M. Kleiner, D. H. Brainard, D. Pelli, What’s new in Psychtoolbox-3? in Perception 36 ECVP Abstract Supplement, (2007).

68. B. F. Rodriguez, S. E. Bruce, M. E. Pagano, M. a Spencer, B. Keller, Factor Structure of the Anxiety Sensitivity Index in a Longitudinal Study of Anxiety Disorder Patients. 42, 79–91 (2004).

69. M. Reuter, H. D. Rosas, B. Fischl, Highly Accurate Inverse Consistent Registration: A Robust Approach. NeuroImage 53, 1181–1196 (2010).

70. M. Reuter, N. J. Schmansky, H. D. Rosas, B. Fischl, Within-Subject Template Estimation for Unbiased Longitudinal Image Analysis. NeuroImage 61, 1402–1418 (2012).

71. A. R. Dykstra, et al., Individualized localization and cortical surface-based registration of intracranial electrodes. 59, 3563–3570 (2012).

72. N. Peled, et al., Invasive Electrodes Identification and Labeling. GitHub Repos. https://gi (2017).

73. B. Fischl, et al., Automatically Parcellating the Human Cerebral Cortex. Cereb. Cortex 14, 11–22 (2004).

74. I. Basu, et al., Consistent Linear and Non-Linear Responses to Electrical Brain Stimulation Across Individuals and Primate Species. Brain Stimulat. 12, 877–892 (2019).

