## Supplemental Figures and Tables for "Bidirectional modulation of human emotional conflict resolution using intracranial stimulation"

**This PDF file includes:**

Supplemental Figures 1-7

Supplemental Tables 1-4

**Supplemental Table 1. Summary table of diagnostic details**. BD-bipolar disorder; GAD-generalized anxiety disorder; MDD- Major Depressive Disorder; PTSD- post-traumatic stress disorder; OCD- obsessive-compulsive disorder.

| *Participant designation* | *Completed psychometric questionnaires* | *Diagnoses* |
|---|---|---|
| *EP01* | | Epilepsy |
| *EP02* | Yes | Epilepsy |
| *EP03* | | Epilepsy |
| *EP04* | | Epilepsy |
| *EP05* | | Epilepsy |
| *EP06* | Yes | Epilepsy |
| *EP07* | Yes | Epilepsy (MDD, past episode) |
| *EP08* | Yes | Epilepsy |
| *EP09* | Yes | Epilepsy |
| *EP10* | Yes | Epilepsy |
| *EP11* | Yes | Epilepsy |
| *EP12* | Yes | Epilepsy |
| *EP13* | Yes | Epilepsy |
| *EP14* | Yes | Epilepsy |
| *EP15* | Yes | Epilepsy |
| *EP16* | | Epilepsy |
| *EP17* | | Epilepsy |
| *EP18* | Yes | Epilepsy, MDD |
| *EP19* | Yes | Epilepsy |
| *EP20* | Yes | Epilepsy |
| *EP21* | Yes | Epilepsy |
| *EP22* | Yes | Epilepsy |
| *EP23* | Yes | Epilepsy |
| *EP24* | Yes | Epilepsy |
| *EP25* | Yes | Epilepsy |
| *EP26* | Yes | Epilepsy |
| *EP27* | Yes | Epilepsy |
| *EP28* | Yes | Epilepsy |
| *EP29* | Yes | Epilepsy |
| *EP30* | Yes | Epilepsy |
| *EP31* | | Epilepsy |
| *EP32* | Yes | Epilepsy |
| *EP33* | | Epilepsy |
| *EP34* | Yes | Epilepsy |
| *EP35* | | Epilepsy |
| *EP36* | | Epilepsy |
| *EP37* | | Epilepsy |
| *EP38* | | Epilepsy |

| | | |
|---|---|---|
| *EP39* | | Epilepsy |
| *EP40* | | Epilepsy |
| *EP41* | | Epilepsy |
| *PP01* | Yes | BD-II |
| *PP02* | Yes | BD (mixed), PTSD, BN |
| *PP03* | Yes | MDD, GAD |
| *PP04* | Yes | BD |
| *PP05* | Yes | MDD, GAD |
| *PP06* | Yes | GAD, panic agoraphobia, recurrent depression |
| *PP07* | Yes | GAD |
| *PP08* | Yes | GAD, panic agoraphobia, past depression |
| *PP09* | Yes | PTSD, MDD, GAD, social phobia |
| *PP10* | Yes | GAD |
| *PP11* | Yes | MDD, GAD, Panic Disorder, Social phobia, PTSD, trichotillomania |
| *PP12* | Yes | BD-II, panic agoraphobia, PTSD, GAD, past alcohol dependence, borderline personality disorder |
| *PP13* | Yes | recurrent MDD, GAD |
| *PP14* | Yes | OCD, past MDD |
| *PP15* | Yes | MDD recurrent, PTSD, GAD, past panic disorder, past social phobia |
| *PP16* | Yes | MDD recurrent, GAD |

**Supplemental Table 2. Average and standard deviation raw psychometric scores per patient group**

| *Participant group* | *DERS Goals* | *ERS* | *ASI* |
|---|---|---|---|
| *Healthy control group* | 11.86 ± 0.602 | 14.68 ± 1.972 | 12.08 ± 1.141 |
| *Patients with psychiatric diagnoses* | 17.20 ± 1.191 | 43.11± 4.747 | 22.55 ± 4.021 |
| *Patients with intractable epilepsy* | 13.07 ± 0.740 | 28.00 ± 2.643 | 16.83 ± 2.252 |

**Supplemental Table 3. Summary table of all ECR with stimulation experiments.** L indicates left hemisphere. R is right hemisphere stimulation. '#b' indicates the number of ~60 trial blocks performed with intermittent stimulation. EP## are identifications per participant. Abbreviations: Dorsolateral (dl) and dorsomedial (dm) prefrontal cortex- dlPFC, dmPFC; lateral (l) and medial (m) orbitofrontal cortex – lOFC, mOFC; ventrolateral prefrontal cortex- vlPFC; dorsal (d) and rostral (r) anterior cingulate- dACC, rACC. BD-bilateral depths; UD-unilateral depths; GrD-Grids and depth electrodes.

| ***Participant designation*** | ***Performed ECR task in the EMU*** | ***Performed ECR task with MEG*** | ***Completed psychometric questionnaires*** | ***Stimulation brain location during the ECR task*** |
|---|---|---|---|---|
| *EP01* | | Yes | | |
| *EP02* | | Yes | Yes | |
| *EP03* | Yes | Yes | | |
| *EP04* | | Yes | | |
| *EP05* | Yes | | | Amygdala, dACC |
| *EP06* | Yes | | Yes | Amygdala, dmPFC |
| *EP07* | | Yes | Yes | |
| *EP08* | Yes | | Yes | Amygdala, rACC, dmPFC |
| *EP09* | Yes | | Yes | dACC, rACC |
| *EP10* | Yes | | Yes | rACC |
| *EP11* | Yes | | Yes | dlPFC, rACC, dACC |
| *EP12* | Yes | Yes | Yes | Amygdala, rACC |
| *EP13* | Yes | Yes | Yes | dlPFC, rACC, dACC |
| *EP14* | | Yes | Yes | |
| *EP15* | Yes | Yes | Yes | dlPFC, dACC, dmPFC |
| *EP16* | | Yes | | |
| *EP17* | Yes | | | dlPFC, dACC |
| *EP18* | Yes | Yes | Yes | |
| *EP19* | Yes | | Yes | Amygdala, dlPFC, dACC |
| *EP20* | Yes | | Yes | dmPFC, dACC |
| *EP21* | Yes | | Yes | dmPFC, dACC |
| *EP22* | Yes | | Yes | |
| *EP23* | Yes | Yes | Yes | dmPFC, dACC |
| *EP24* | Yes | Yes | Yes | dmPFC, dACC |
| *EP25* | Yes | | Yes | |
| *EP26* | Yes | | Yes | |
| *EP27* | Yes | | Yes | |
| *EP28* | Yes | | Yes | |
| *EP29* | Yes | Yes | Yes | dmPFC, dACC |
| *EP30* | Yes | Yes | Yes | |
| *EP31* | Yes | | | |

| | | | |
|---|---|---|---|
| *EP32* | Yes | | Yes |
| *EP33* | Yes | | |
| *EP34* | Yes | | Yes |
| *EP35* | Yes | | |
| *EP36* | Yes | | |
| *EP37* | Yes | | |
| *EP38* | Yes | | |
| *EP39* | Yes | | |
| *EP40* | Yes | | |
| *EP41* | Yes | | |

**Supplemental Table 4. Statistical comparisons of behavioral changes between stimulation sites, including the dlPFC, dmPFC, rACC, dACC, and amygdala.** State estimate variables described in **Supplemental Figure 3**.

| ***Behavioral measure*** | ***Chi-square, Kruskal-Wallis test*** | ***p- value, Kruskal-Wallis test*** |
|---|---|---|
| *Reaction time, during stimulation trials* | 16.6819 | 0.0022 |
| *Reaction time, during non-stimulation trials* | 6.7149 | 0.1517 |
| *Model 1, RTB* | 4.948034 | 0.422255 |
| *Model 1, ECS* | 4.240179 | 0.51538 |
| *Model 1, EAS* | 7.212946 | 0.205279 |
| *Model 2, RTB* | 5.462277 | 0.3621 |
| *Model 2, CvsI* | 4.707533 | 0.452605 |
| *Model 2, Tr.* | 8.536775 | 0.129029 |
| *Model 4, RTB* | 4.981715 | 0.418116 |
| *Model 4, Tr.* | 9.493049 | 0.090942 |
| *Model 10, RTB* | 6.314974 | 0.276766 |
| *Model 10, CvsI* | 3.102176 | 0.684237 |
| *Model 10, CC* | 5.345337 | 0.375203 |
| *Model 12, RTB* | 5.038411 | 0.41121 |
| *Model 12, Tr.* | 9.318601 | 0.097011 |
| *Model 15, RTB* | 4.62549 | 0.463271 |
| *Model 15, CvsI* | 5.726904 | 0.333706 |
| ***Model 15, Conflict-Adaptation*** | **14.72865** | **0.011587** |
| *Model 19, RTB* | 4.793006 | 0.441661 |
| *Model 19, CvsI* | 0.291859 | 0.997793 |
| *Model 19, Valence* | 4.37487 | 0.496798 |
| *Model 21, RTB* | 4.998611 | 0.41605 |
| *Model 21, Face Valence* | 3.428689 | 0.634204 |
| *Model 21, Word Valence* | 0.294761 | 0.99774 |

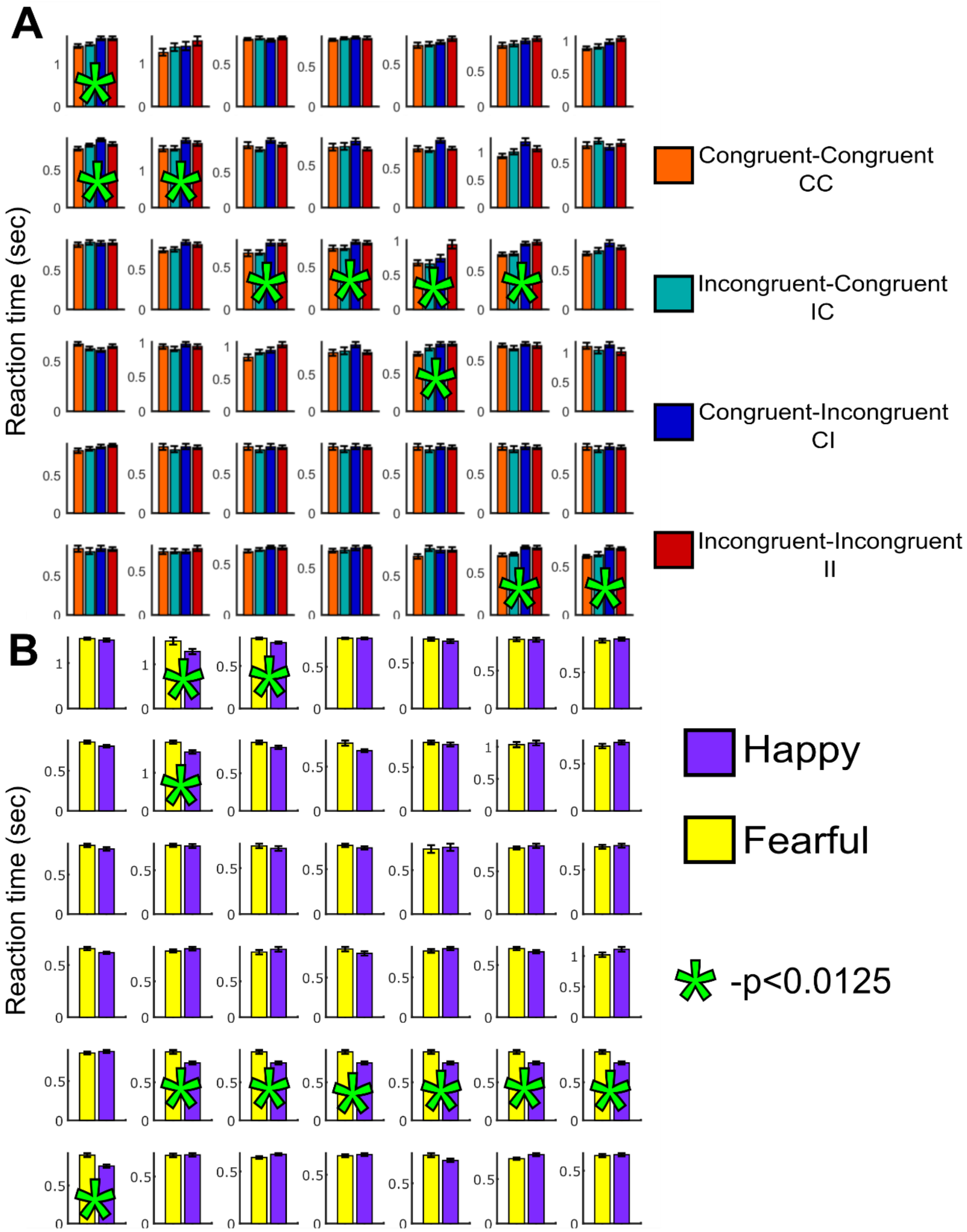

**Supplemental Figure 1. Participants respond differently to ECR task features while the electrode locations could be mapped to similar areas across the brain**. **A**. Behavioral differences between trial types on a per-participant level (N=41, two figures are repeated tasks within the same individual). In the task, participants are asked to identify the emotion on the face while ignoring the word. The face and the word can match (congruent (C) trials) or conflict (incongruent (I) trials; Etkin *et al.*, 2006). The changes in trial types (e.g. congruent to incongruent, or CI) as well as trial types induce differences in reaction times across the trial types across multiple, but not all, participants (* indicates significant differences between trial types, $p<0.0125$, Kruskal-Wallis test). **B**. There were no significant differences between happy versus fearful face trials (* indicates significant differences between trial types, p<0. 0125, Wilcoxon rank sum test). Error bars indicate standard error across trials per participant.

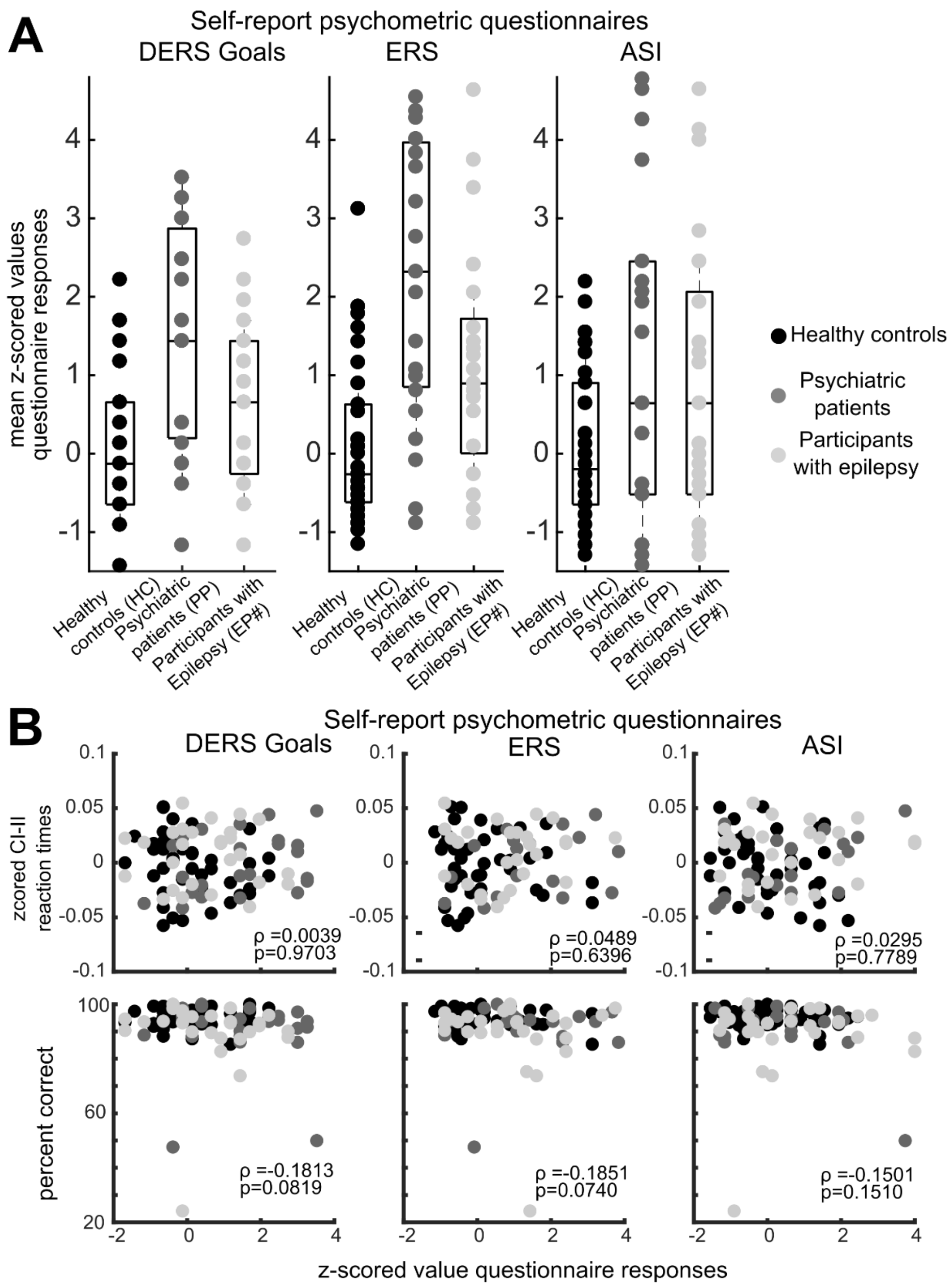

**Supplemental Figure 2. A.** z-scored participant scores in response to three psychometric questionnaires, namely Emotional Reactivity Scale (ERS), Difficulties in Emotion Regulation Scale (DERS), and Anxiety Sensitivity Index (ASI). ERS measures emotion reactivity, DERS measures emotion regulation skills, and ASI sensitivity to anxiety-related physiological symptoms. **B.** Spearman correlation between zscored psychometric questionnaire scores and the difference in z-scored reaction times during CI vs II trials (top) and the correlation between the zscored psychometric questionnaire scores and the % correct (bottom) per psychometric scale.

### Different state estimate models

##### Models testing how trial history and congruence can describe and predict behavior

**Model 1: RTB, II, CI**

$\log Y_k = X_{k,1} + X_{k,2} * CI_k + X_{k,3} * II_k + v_k \quad v_k \sim N(0, {\sigma_v}^2)$

**Model 15: RTB, CvsI,CI (also CI-II from Model 1)**

$\log Y_k = X_{k,1} + X_{k,2} * I_k + X_{k,3} * CI_k + v_k \quad v_k \sim N(0, {\sigma_v}^2)$

**Model 2 : RTB, CvsI, Tr.**

$\log Y_k = X_{k,1} + X_{k,2} * CvsI_k + X_{k,3} * Tr_k + v_k \quad v_k \sim N(0, {\sigma_v}^2)$

**Model 3 : RTB, CvsI**

$\log Y_k = X_{k,1} + X_{k,2} * CvsI_k + v_k \quad v_k \sim N(0, {\sigma_v}^2)$

**Model 4: RTB and Tr.**

$\log Y_k = X_{k,1} + X_{k,2} * Tr_k + v_k \quad v_k \sim N(0, {\sigma_v}^2)$

**Model 5: CvsI, Tr.**

$\log Y_k = X_{k,2} * CvsI_k + X_{k,3} * Tr_k + v_k \quad v_k \sim N(0, {\sigma_v}^2)$

**Model 18: RTB**

$\log Y_k = X_{k,1} + v_k \quad v_k \sim N(0, {\sigma_v}^2)$

**Model 7: RTB, CI, IC**

$\log Y_k = X_{k,1} + X_{k,2} * CI_k + X_{k,3} * IC_k + v_k \quad v_k \sim N(0, {\sigma_v}^2)$

**Model 10: RTB, CvsI, CC**

$\log Y_k = X_{k,1} + X_{k,2} * CvsI_k + X_{k,3} * CC_k + v_k \quad v_k \sim N(0, {\sigma_v}^2)$

**Model 16: RTB, CvsI,CI, trial number**

$\log Y_k = X_{k,1} + X_{k,2} * I_k + X_{k,3} * CI_k + X_{k,3} * trial_k + v_k \quad v_k \sim N(0, {\sigma_v}^2)$

##### Models testing how trial valence can describe and predict behavior

**Model 19: RTB, CvsI, Face Valence)**

$\log Y_k = X_{k,1} + X_{k,2} * I_k + X_{k,3} * CI_k + X_{k,3} * trial_k + v_k \quad v_k \sim N(0, {\sigma_v}^2)$

**Model 20: RTB, Face Valence)**

$\log Y_k = X_{k,1} + \beta_1 * ValenceFace_k + \beta_2 * Tr_k + v_k \quad v_k \sim N(0, {\sigma_v}^2)$

**Model 21: RTB, Face Valence, Word Valence):**

$\log Y_k = X_{k,1} + \beta_1 * ValenceFace_k + \beta_2 * ValenceWord_k + v_k \quad v_k \sim N(0, {\sigma_v}^2)$

**Model 22: RTB, Word Valence)**

$\log Y_k = X_{k,1} + \beta_1 * ValenceWord_k + v_k \quad v_k \sim N(0, {\sigma_v}^2)$

##### Models testing whether fixing some model features can describe and predict the behavior

**Model 11: RTB, CI, IC, fix CvsI**

$\log Y_k = X_{k,1} + X_{k,2} * CI_k + X_{k,3} * IC_k + \beta_1 * CvsI_k + v_k \quad v_k \sim N(0, {\sigma_v}^2)$

**Model 6: CI, IC, fix CvsI, and RTB**

$\log Y_k = \beta_0 + X_{k,2} * CI_k + X_{k,3} * IC_k + \beta_1 * CvsI_k + v_k \quad v_k \sim N(0, {\sigma_v}^2)$

**Model 8: CI, IC, fix RTB**

$\log Y_k = \beta_0 + X_{k,2} * CI_k + X_{k,3} * IC_k + v_k \quad v_k \sim N(0, {\sigma_v}^2)$

**Model 9: CC, CI, fix RTB**

$\log Y_k = X_{k,1} + X_{k,2} * CC_k + X_{k,3} * CI_k + v_k \quad v_k \sim N(0, {\sigma_v}^2)$

**Model 12: RTB, Tr., fix CvsI**

$\log Y_k = X_{k,1} + X_{k,2} * Tr_k + \beta_1 * CvsI_k + v_k \quad v_k \sim N(0, {\sigma_v}^2)$

**Model 13: RTB, fix CvsI**

$\log Y_k = X_{k,1} + \beta_1 * CvsI_k + v_k \quad v_k \sim N(0, {\sigma_v}^2)$

**Model 14: RTB, fix CvsI and Tr. and intercept**

$\log Y_k = X_{k,1} + \beta_1 * CvsI_k + \beta_2 * Tr_k + \beta_3 + v_k \quad v_k \sim N(0, {\sigma_v}^2)$

**Model 17: RTB, fix CvsI and Tr.**

$\log Y_k = X_{k,1} + \beta_1 * CvsI_k + \beta_2 * Tr_k + v_k \quad v_k \sim N(0, {\sigma_v}^2)$

##### Decision models using only trial accuracy to model states:

**Model 23: RTB, II, CI**

$\mathrm{logit}(p(d_k = 1)) = X_{k,1} + X_{k,2} * CI_k + X_{k,3} * I2I_k$

$d_k \in \{'True','False'\} \equiv \{1,0\}$

**Model 24: RTB, CvsI, Tr.**

$\mathrm{logit}(p(d_k = 1)) = X_{k,1} + X_{k,2} * CvsI_k + X_{k,3} * Tr_k$

$d_k \in \{'True','False'\} \equiv \{1,0\}$

**Model 25: RTB, CvsI,CI (also CI-II from Model 1)**

$\mathrm{logit}(p(d_k = 1)) = X_{k,1} + X_{k,2} * I_k + X_{k,3} * CI_k$

$d_k \in \{'True','False'\} \equiv \{1,0\}$

**Model 26: RTB and Tr.**

$\mathrm{logit}(p(d_k = 1)) = X_{k,1} + X_{k,2} * Tr_k$

$d_k \in \{'True','False'\} \equiv \{1,0\}$

##### Mixed models using both reaction time and trial accuracy to model states:

**Model 27: RTB, II, CI**

$\log Y_k = X_{k,1} + X_{k,2} * CI_k + X_{k,3} * II_k + v_k \quad v_k \sim N(0, {\sigma_v}^2)$

$\mathrm{logit}(p(d_k = 1)) = \beta_0 + \beta_1 * X_{k,1} + \beta_2 * X_{k,2} * CI_k + \beta_3 * X_{k,3} * II_k$

$d_k \in \{'True','False'\} \equiv \{1,0\}$

**Model 28: RTB, CvsI, Tr.**

$\log Y_k = X_{k,1} + X_{k,2} * CvsI_k + X_{k,3} * Tr_k + v_k \quad v_k \sim N(0, {\sigma_v}^2)$

$\mathrm{logit}(p(d_k = 1)) = \beta_0 + \beta_1 * X_{k,1} + \beta_2 * X_{k,2} * CvsI_k + \beta_3 * X_{k,3} * Tr_k$

$d_k \in \{'True','False'\} \equiv \{1,0\}$

**Model 29: RTB, CvsI,CI (also CI-II from Model 1)**

$\log Y_k = X_{k,1} + X_{k,2} * I_k + X_{k,3} * CI_k + v_k \quad v_k \sim N(0, {\sigma_v}^2)$

$\mathrm{logit}(p(d_k = 1)) = \beta_0 + \beta_1 * X_{k,1} + \beta_2 * X_{k,2} * I_k + \beta_3 * X_{k,3} * CI_k$

$d_k \in \{'True','False'\} \equiv \{1,0\}$

**Model 30: RTB and Tr.**

$\log Y_k = X_{k,1} + X_{k,2} * Tr_k + v_k \quad v_k \sim N(0, {\sigma_v}^2)$

$\mathrm{logit}(p(d_k = 1)) = \beta_0 + \beta_1 * X_{k,1} + \beta_2 * X_{k,2} * Tr_k$

$d_k \in \{'True','False'\} \equiv \{1,0\}$

##### state estimate terms

RTB: reaction time bias state

CC: congruent-congruent trial state

CI: congruent-incongruent trial state

Tr.: Transition trial state, e.g. IC or CI

II: incongruent-incongruent trial state

IC: incongruent-congruent trial state

CvsI: Congruent or Incongruent trial state

Face Valence: valence (happy-fear) of the face state

Word Valence: valence (happy-fear) of the word state

trial number: trial number in the task

fix: Fixed state estimate value across trials

**Supplemental Figure 3. Developing and testing ECR state estimate models on the behavior.** The models tested use similar approaches to Model 1 (**Fig. 2A**) though the indicator terms and what terms were fixed varied from model to model and are laid out per model here. The legend below indicate the task-relevant information for the terms in the model. 'Fix' descriptors for each model indicates that the term was fixed in the model and was not estimated as a state through the expectation maximization algorithm. The models could be subdivided into three main groups: 1) models which include terms relating to trial history or congruence; 2) models which include valence such as happy or fearful stimuli; 3) Models which have 'fixed' terms which do not vary in the course of the model estimation.

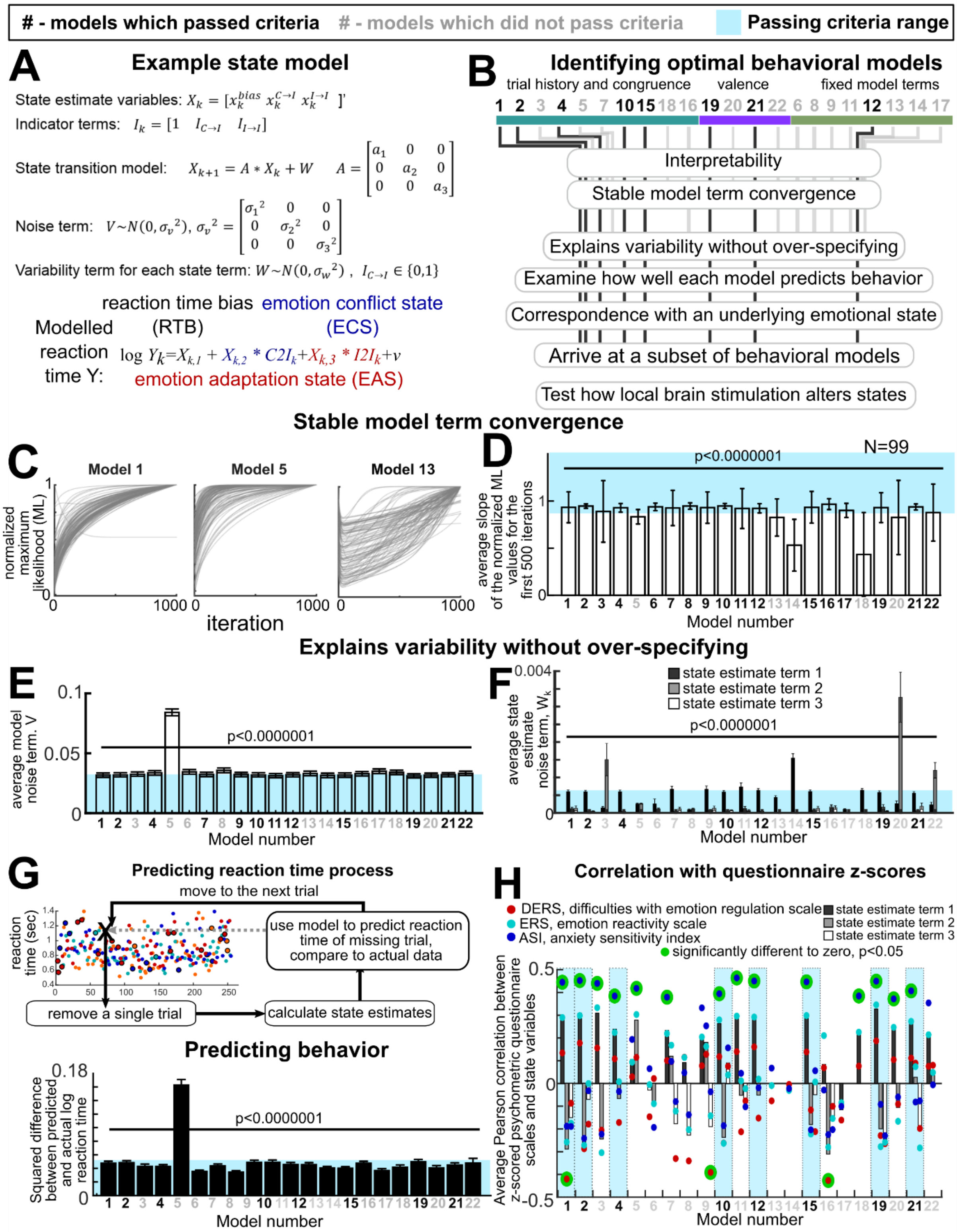

**Supplemental Figure 4**. **Developing and testing ECR state estimate models of behavior using only non-stimulated task sessions. A**. Example state estimate model: ECR state estimate

Model 1. The ECR state estimation is broken into three main components: the overall reaction time state (Bias), the emotion conflict state (C2I), and the adaptation state (I2I). The noise terms in the transition state (V) as well as the noise terms pertaining to each state term (W) can be used to identify which models are optimal for the ECR behavioral data (C). **B**. The protocol for identifying the ideal model for the ECR state estimate involved testing 30 models on the behavior of multiple participant groups using multiple layers of criteria. Model numbers indicated in bold black print were identified as viable while model numbers in grey were not. The twenty-two models could be subdivided into three main groups: 1) models which include terms relating to trial history or congruence (indicated by a teal bar); 2) models which include valence such as happy or fearful stimuli (indicated by a purple bar); 3) models which have ‘fixed’ terms which do not vary in the course of the model estimation and was not estimated as a state (indicated by a green bar). Only 22 of the 30 total mdels are shown here. **C.** Example maximum likelihood estimates over 1000 model iterations. **D**. Each model displayed different maximum likelihood slopes on the same ECR behavioral data (N=99). For the same non-stimulated task session behavioral data (N=99), we compared the overall noise term ( **E** ), the state variable noise term ($W_k$, **F**), and the RMS between the actual and predicted reaction times using a leave one out strategy (**G**). **H**. The Pearson correlations between z-scored individual psychometric questionnaire scores and the state variable, only using non-stimulated task ECR behavioral data (N=99). Labels for each bar for the state estimates are indicated in the **F**. In **D-G**: blue areas indicate threshold ranges used to pass or reject models based on the plotted terms. p-value indicates significant differences between terms per model and state variable, Friedman test, $p<0.0001$.

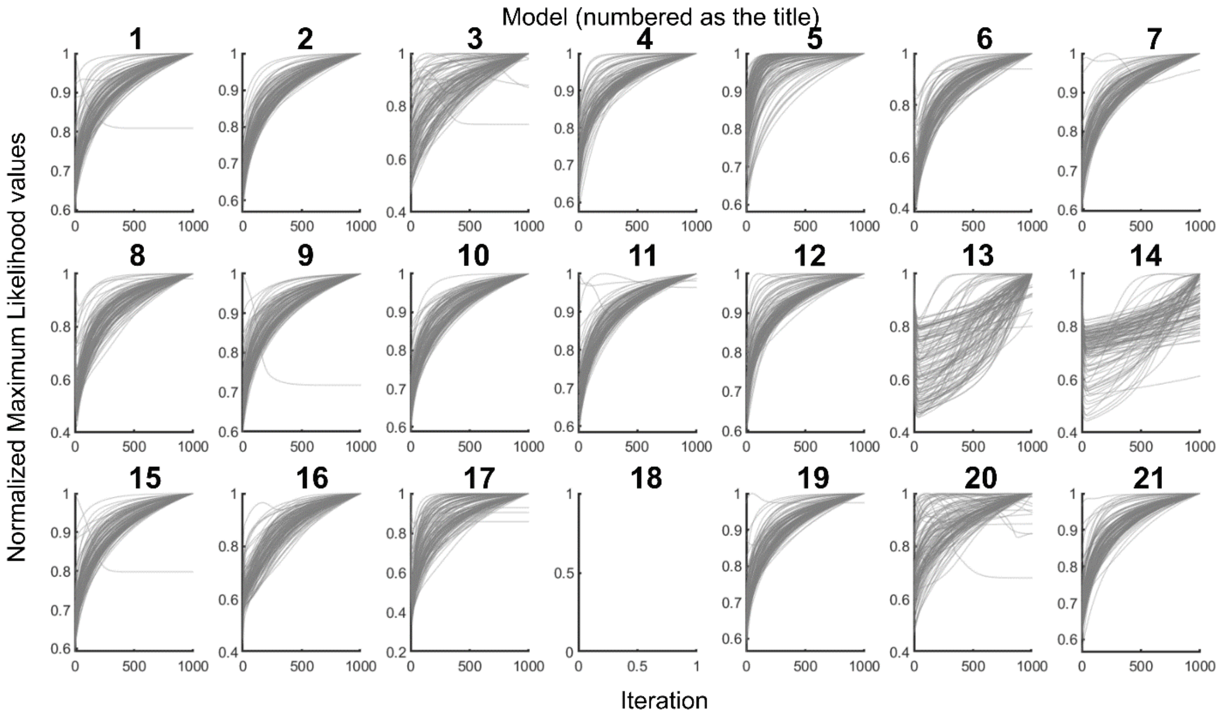

**Supplemental Figure 5. Normalized maximum likelihood over 1000 iterations.**

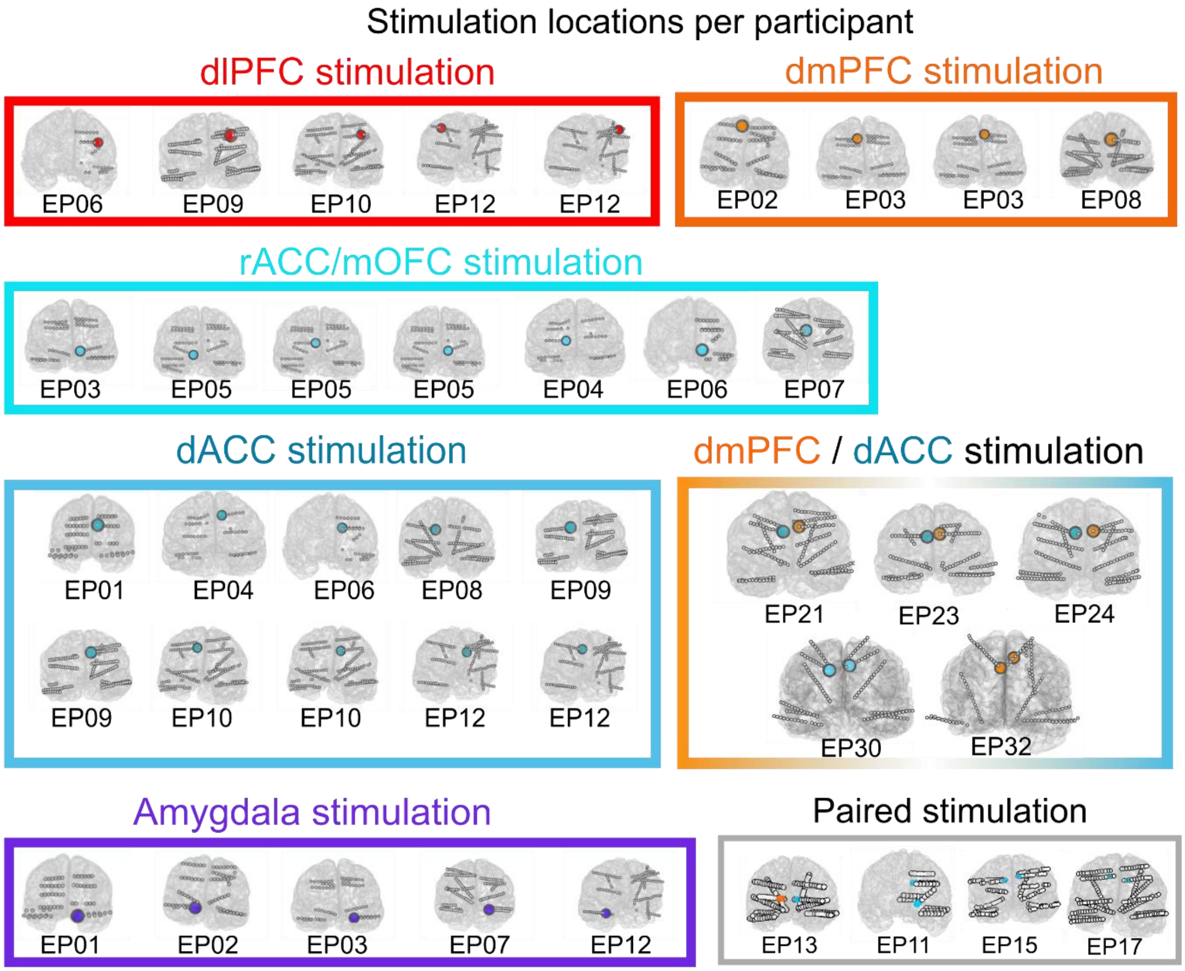

**Supplemental Figure 6. Nineteen epilepsy participants have been stimulated while performing the ECR task across multiple brain regions. A**. In this summary figure of all the electrode sites where the participants were stimulated with either 130 or 160 Hz frequencies for 400 msec at the image onset. dmPFC/dACC stimulation indicate stimulation of both areas intermittently in the same block. Abbreviations: Dorsolateral (dl) and dorsomedial (dm) prefrontal cortex- dlPFC, dmPFC; lateral (l) and medial (m) orbitofrontal cortex – lOFC, mOFC; ventrolateral prefrontal cortex- vlPFC; dorsal (d) and rostral (r) anterior cingulate- dACC, rACC.

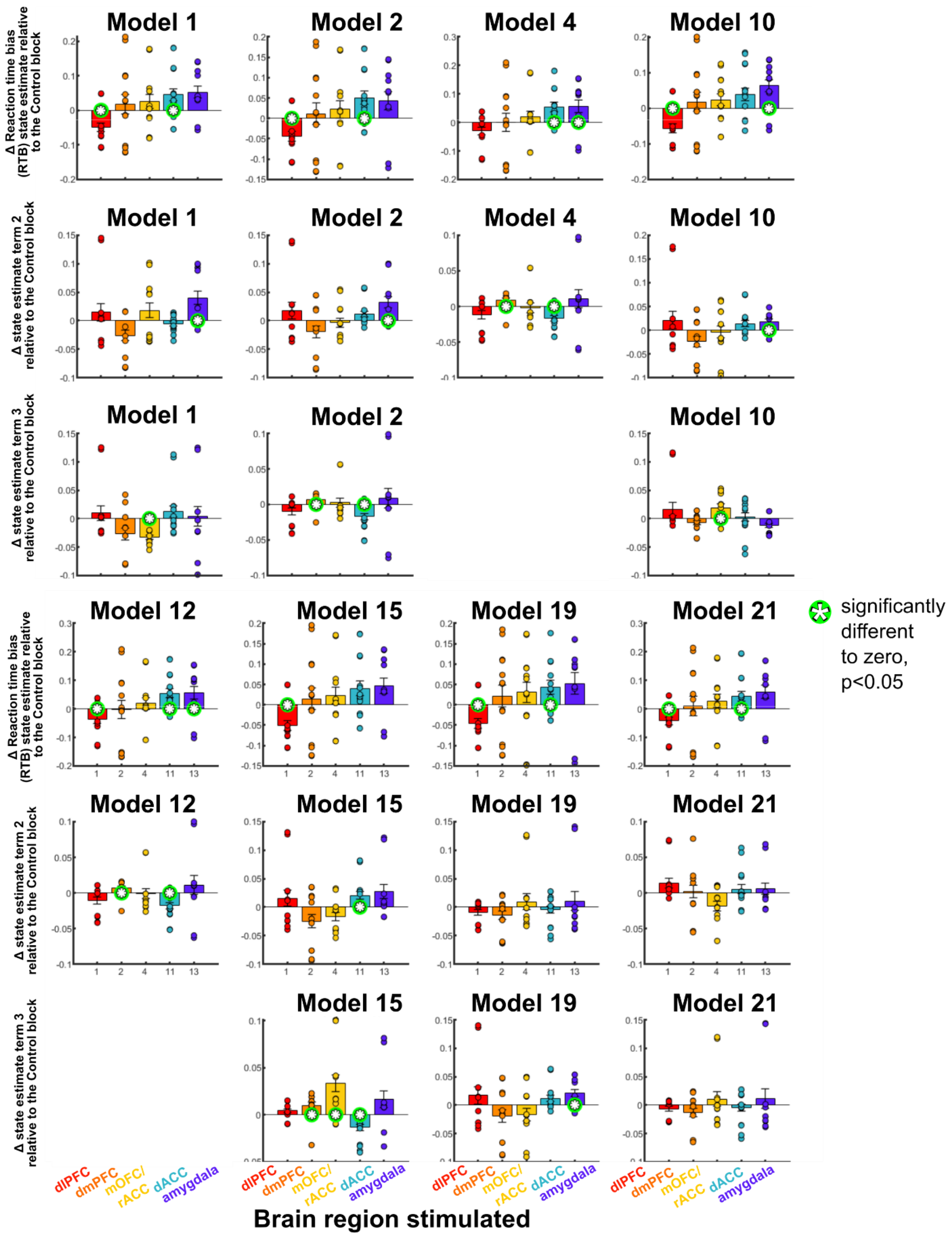
Model 1
Model 2
Model 4
Model 10
Model 12
Model 15
Model 19
Model 21
Δ Reaction time bias (RTB) state estimate relative to the Control block
Δ state estimate term 2 relative to the Control block
Δ state estimate term 3 relative to the Control block
significantly different to zero, p<0.05
dlPFC
dmPFC
mOFC/ rACC
dACC
amygdala
Brain region stimulated

**Supplemental Figure 7**. **Neural stimulation alters the state estimate dynamics**. Five models have been tested on the behavior of epilepsy participants performing the ECR task during stimulation (N=13). Average changes in state estimate values with stimulation across epilepsy participants. We compared the Test blocks (interleaved stimulation trials) to the Control blocks (no stimulation). Changes in average state estimates across participants per stimulation site, color coded in the bars. The error bars indicate standard error across participants (N=13). State estimates are indicated along the y axis for each model. The top row is the bias term while the lower rows are the other state estimate terms (e.g. CI). Figures without p-values are not significantly different between stimulation site conditions. Red circles at the bar (for each brain region) indicate which changes from the Control blocks during Test blocks are significantly different from zero, Wilcoxon sign-rank test, $p<0.05$. Brain region abbreviations as indicated in **Supplemental Figure 6**.
